# Inhibitor of the spindle assembly checkpoint surpasses apoptosis sensitizer in synergy with taxanes

**DOI:** 10.1101/414359

**Authors:** Teng-Long Han, Zhi-Xin Jiang, Yun-Tian Li, Deng-Shan Wu, Jun Ji, Hu Lin, Bin Hu, Hang Sha

## Abstract

The antitumor effect of taxanes have been attributed to their ability to induce mitotic arrest through activation of the spindle assembly checkpoint. Cell death following prolonged mitotic arrest is mediated by the intrinsic apoptosis pathway. Thus, apoptosis sensitizers which inhibit antiapoptotic Bcl-2 family proteins has been shown to enhance taxanes-induced cell death. By contrast, spindle checkpoint disruption facilitates mitotic slippage and is thought to promote taxanes resistance. Notably, other modes of cell death also contribute to treatment outcomes. Here we show that inhibition of the spindle checkpoint suppresses taxanes induced apoptosis but increases terminal growth arrest of tumor cells with features of cellular senescence. By using clonogenic assay which measures the net result of multiple forms of cell death and is more reflective of therapeutic response, our finding suggests apoptosis is not a major determinant of antitumor efficacy of taxanes, whereas spindle checkpoint inhibitor displays a long-term advantage over apoptosis sensitizer in blocking colony outgrowth of tumor cells when combined with different microtubule toxins, therefore represents a superior therapeutic strategy.

**SIGNIFICANCE:** Apoptosis has long been regarded as the primary mechanism of anti-cancer efficacy of taxanes, while the role of the spindle assembly checkpoint (SAC) in treatment response to taxanes has been controversial. Either apoptosis sensitizer or inhibitor of SAC has been reported to synergize with taxanes. While inhibitor of antiapoptotic proteins potentiates taxanes induced apoptosis, inhibitor of SAC suppresses apoptosis by facilitating mitotic slippage, that is why it is implicated in taxanes resistance. By demonstrating that apoptotic rates are not associate with long-term treatment response, not only do we find that inhibitor of SAC displays a long-term advantage over apoptosis sensitizer in combination with taxanes, but we also resolve the dispute around the role of SAC in cellular response to taxanes.

## INTRODUCTION

Docetaxel and paclitaxel are taxanes widely used in clinic for the treatment of ovarian, breast, lung cancer and other solid tumors (Montero et al., 2005). Though docetaxel appears to be more potent than paclitaxel (Verweij et al., 1994), they share a common mechanism of action by promoting and stabilizing microtubule assembly, which disrupt microtubule dynamics during mitosis and lead to chronic activation of the spindle assembly checkpoint (SAC) and mitotic arrest of tumor cells (Rieder and Maiato, 2004). Prolonged mitotic arrest followed by either death during mitosis or an abnormal exit from mitosis, to a tetraploid G1 state known as mitotic slippage. After slippage, cells may die, arrest, or continue to proliferate (Weaver and Cleveland, 2005). Some have hypothesized that the length of mitotic arrest determines whether a cell die during mitotic arrest, as well as its fate after mitotic exit, with cells that arrest longer being more likely to die, though this issue is still controversial (Weaver, 2014).

Taxanes’ antitumor effect has been attributed to their ability to induce mitotic arrest. Although the molecular link between mitotic arrest and cell death remains unclear, the accumulation of proapoptotic signals during prolonged mitotic arrest has been observed by many groups (Gascoigne and Taylor, 2008; Huang et al., 2009). Cell death during mitotic arrest or after mitotic exit is thought to occur through the intrinsic apoptosis pathway (Shi et al., 2011), the primary mechanism for cell death following many types of chemotherapy. Accordingly, disruption of apoptotic responses seems to be a major contributor to treatment resistance (Adams and Cory, 2007). For example, overexpression of antiapoptotic members of the Bcl-2 family, the central regulator of intrinsic apoptosis pathway, are frequently associated with resistance of many tumors to chemotherapy (Amundson et al., 2000). Therefore, targeting the antiapoptotic Bcl-2 family members offers an attractive opportunity to reverse chemotherapeutic drug resistance. Recently, small molecule inhibitors which mimic BH3-activity by binding with high affinity to antiapoptotic Bcl-2 family members have been developed. These ‘BH3-mimetics’ (i.e., ABT-737, ABT-263 and ABT-199) displays preclinical activity as single agent, and acts synergistically with chemotherapeutic agents such as taxanes, thus hold big promise for anti-cancer therapy (Vogler et al., 2009). Of note, ABT-199 (venetoclax) was recently approved by the US Food and Drug Administration (FDA) for the treatment of chronic lymphocytic leukemia (CLL) with a specific chromosomal abnormality (Roberts et al., 2016).

Mitotic arrest induced by taxanes depends on an active SAC, a key surveillance mechanism that delays anaphase onset until all chromosomes are properly attached to a bipolar mitotic spindle, thus ensuring an accurate segregation of chromosomes during mitosis. The role of SAC in cellular response to taxanes has been controversial. Although complete inhibition of SAC is lethal to cell (Kops et al., 2004) and targeting SAC was reported to enhances the cytotoxic effects of taxanes, an effect that has been attributed to intolerable enhancement of chromosome mis-segregation (Janssen et al., 2009), cells with a weakened checkpoint survive and exhibit chromosomal instability (CIN) and aneuploidy (Weaver and Cleveland, 2005), both phenotypes are commonly found in cancer cells and are thought to contribute to intrinsic resistance to taxanes (Swanton et al., 2007; Swanton et al., 2009). Moreover, a compromised SAC facilitates mitotic exit thus enable tumor cells to escape apoptosis that otherwise occurs during prolonged mitotic arrest following paclitaxel treatment. Accordingly, a functional SAC has been reported by numerous groups to be required for efficient cell killing by paclitaxel (Sudo et al., 2004; Tao et al., 2005).

In an attempt to clarify the role of SAC in cellular response to taxanes, we found the type of assays used may well affect the conclusion as to whether inhibition of SAC sensitize tumor cells or confer drug resistance. Many studies choose to draw their conclusions through measuring the amount of cell death or cell viability within a short time frame following treatment (Sudo et al., 2004; Swanton et al., 2007). However, delayed cell death is common following therapy and other less-acute form of cell death (eg. Senescence) also contribute to treatment outcomes, particularly at sub-apoptotic concentrations (Rebbaa et al., 2003), thus cell death could be severely underestimated when assayed by short-term analyses, such as those measuring apoptosis. By using clonogenic assay, which simultaneously measure the net result of multiple forms of death and examine cellular response with longer-term kinetics that are more reflective of therapeutic response (Brown and Attardi, 2005), we find the fraction of apoptosis does not predict treatment outcomes and the combination of SAC inhibition and taxanes displays a clear advantage over the combination with apoptosis sensitizer in blocking the long-term growth of tumor cells.

## Results

### The long-term effect of Docetaxel correlates with drug concentration rather than the duration of mitotic arrest

Although the mechanisms by which taxanes kill tumor cells remain obscure, what is clear is that by targeting and stabilizing microtubule, taxane treatment leads to chronic activation of SAC thereby inducing a prolonged mitotic arrest (Gascoigne and Taylor, 2009). We found the robustness of mitotic arrest depends on drug concentration. SAC proficient HeLa cells (Masuda et al., 2003) were synchronized and treated with different concentrations of Docetaxel. At 64nM, the cells exhibited sustained mitotic arrest followed by substantial cell death within 32 hr. By contrast, at 8nM, the cells were blocked only transiently followed by mitotic slippage as judged by reduced mitotic index after 16 hr (Figure 1A), indicating the mitotic arrest was unable to maintain by the treatment of lower concentration. Moreover, we found the cell killing by Docetaxel was also correlated with drug concentration, with higher concentration of the drug lead to more significant cell death (Figure 1B).

**Figure 1.**
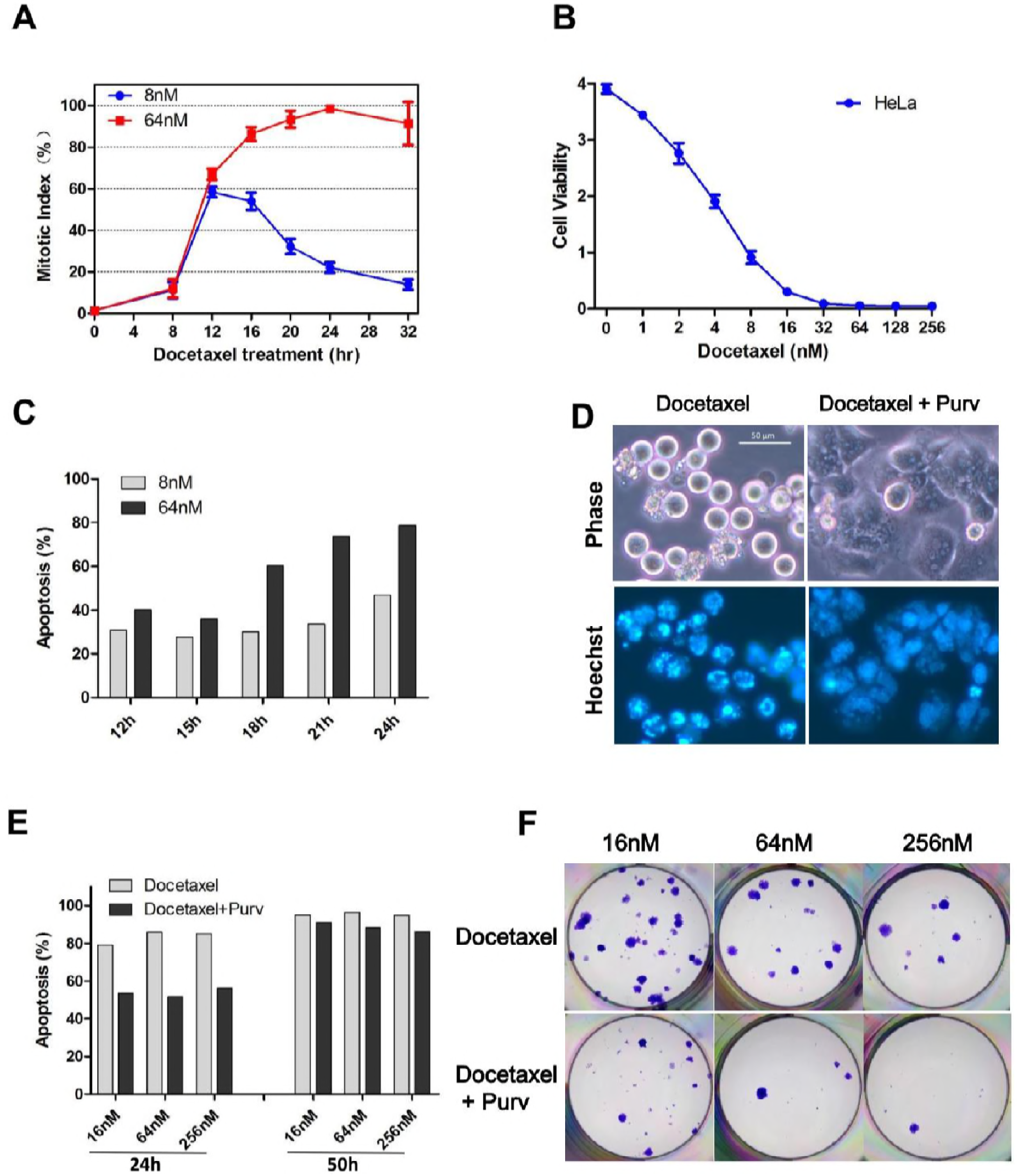
The relationship between mitotic arrest and treatment response to Docetaxel. (A) Synchronized HeLa cells were treated with 8nM or 64nM docetaxel and collected at indicated time intervals. Mitotic indexes were determined by counting the percentage of cells with spherical morphology and condensed chromatin. Values represent the means ± S.D. (n = 3 fields). (B) HeLa Cell viability was determined 3 days after treatment with the various concentrations of docetaxel. Values represent the means ± S.D. (n = 3 wells). (C) HeLa cells were treated as described in (A) and collected at indicated time intervals. Percentage of apoptosis were detected by FACS analysis following Annexin V/PI staining (see Figure S1A for details). (D) Synchronized HeLa cells were exposed to Docetaxel (64nM) for 16 hr and then Purv (10 μM) was added for promotion of mitotic slippage. After 24 hr Docetaxel treatment with (right panels) or without Purv (left panels), the cells were stained with Hoechst 33342 and photographed using phase and Hoechst fluorescence. (E) Synchronized cells were treated different concentrations of Docetaxel followed by the addition of Purv, as described in (D). Cells were collected after 24 hr (left panel) or 50 hr (right panel) Docetaxel treatment with or without Purv, and analyzed for induction of apoptosis (see Figure S1B for details). (F) Colony outgrowth assay of HeLa cells. Cells were treated as described in (E). After 24 hr treatment with Docetaxel in the absence (upper panels) or presence (lower panel) of Purv,the cells were cultured in drug-free medium for 2 weeks and colony formation was photographed following fixation with methanol and staining with crystal violet.

To determine whether cell death induced by higher concentrations was due to prolonged mitotic arrest, induction of Apoptosis was detected at various time points following drug treatment. At 64nM, the fraction of apoptosis increased dramatically following 16 hr Docetaxel treatment, the time point at which the majority of the cells entered and were blocked in mitosis. By contrast, the marked increase of apoptosis was not observed in cells treated with 8nM Docetaxel (Figure 1C and S1A), which only transiently blocked in mitosis. These observations suggest a correlation between prolonged mitotic arrest and mitotic cell death.

To further address this, cells were forced to exit mitosis following 16 hr exposure to docetaxel by addition of purvalanol A (Purv) which inhibit the mitotic kinase Cdk1 thus facilitates mitotic exit despite the activation of SAC, yielding cells in the subsequent interphase with multiple nuclei (Figure 1D right), contrasting with cells with spherical morphology and condensed chromatin that arrest in mitosis (Figure 1D left). The apoptotic cells at 24 hr treatment were markedly reduced by addition of Purv (Figure 1E left and S1B), confirming the contribution of mitotic arrest to cell death. However, similar levels of apoptosis were detected at 50 hr, in the absence or presence of Purv (Figure 1E right and S1B), suggesting that the promotion of mitotic exit by Purv only delayed apoptosis, by switching cell death from mitosis to the subsequent interphase.

Colony formation assays were performed to determine the long-term cellular response to Docetaxel, alone or in combination of Purv. Despite the fact that promotion of slippage by Purv suppressed apoptotic cell death during mitosis, it did not promote long-term cell survival as judged by reduced number of colonies following Purv addition (Figure 1F), though the toxic effect of Purv itself to cell viability cannot be ruled out. Moreover, we found the colony outgrowth following treatment was associated with drug concentration. Higher concentration of Docetaxel always yielded fewer number of colonies, in the absence or presence of Purv. Notably, by promoting mitotic slippage, the addition of Purv unifies the duration of mitotic arrest induced by different concentrations of Docetaxel. Therefore, the superior efficacy of higher concentration of Docetaxel against colony outgrowth should not be attributed to prolonged mitotic arrest.

### Apoptosis rate does not predict colony outgrowth after drug withdraw

Unlike HeLa cells, MDA-MB-231 cells were incapable to sustain a mitotic arrest regardless of drug concentration. Mitotic cell death was not observed in MDA-MB-231 cells. Upon exposure to Docetaxel MDA-MB-231 cells were only blocked transiently followed by mitotic exit (Figure 2A), suggesting a compromised checkpoint response in MDA-MB-231 cells compared with that of HeLa cells.

**Figure 2.**
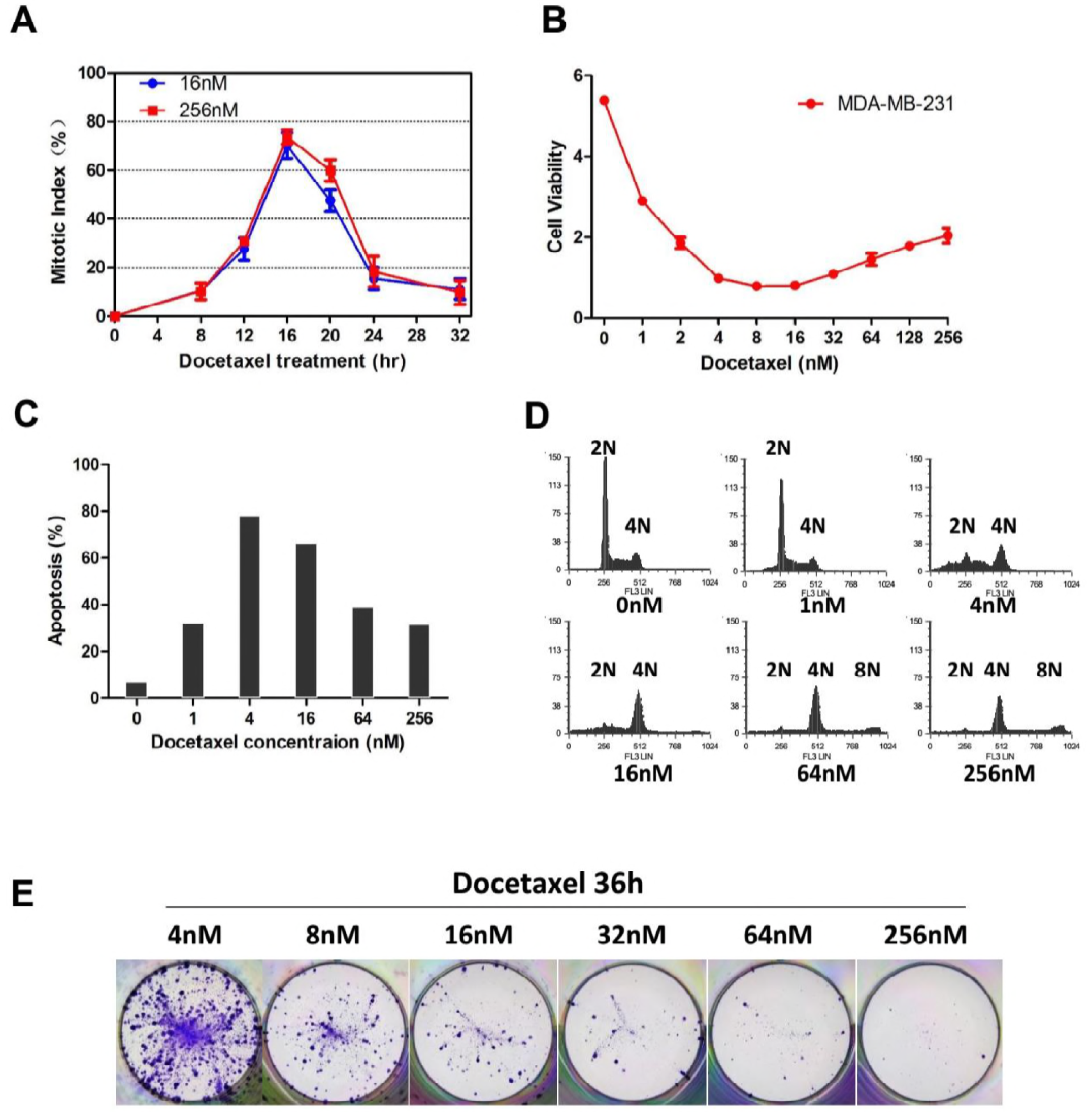
Apoptosis rates does not predict colony outgrowth after drug withdraw. (A) Synchronized MDA-MB-231 cells were treated with different concentration of docetaxel and collected at indicated time intervals. Mitotic indexes were determined by counting the percentage of cells with spherical morphology and condensed chromatin. Values represent the means ± S.D. (n = 3 fields). (B) MDA-MB-231 Cell viability was determined 3 days after treatment with the indicated concentrations of docetaxel. Values represent the means ± S.D. (n = 3 wells). (C and D) MDA-MB-231 cells were treated with the indicated concentrations of docetaxel for 48h, then collected and analyzed for induction of apoptosis (C, see Figure S2A for details) and DNA content (D) by FACS. (E) Cells were treated with the indicated concentrations of docetaxel for 36h, and then cultured in drug-free medium for 4 weeks. Colony outgrowth was photographed following fixation and staining with crystal violet.

We have shown that high concentration of Docetaxel lead to prolonged mitotic arrest in HeLa cells which in turn induced substantial cell death within a short time frame. To examine the cell killing in MDA-MB-231 cells, both Cell viability (Figure 2B) and induction of apoptosis (Figure 2C and S2A) were determined following treatment with increasing concentrations of Docetaxel. Unexpectedly, relatively lower concentrations (4, 16nM) of Docetaxel resulted in more cell death compared with higher concentrations. To ascertain the cause of these different responses, Cell cycle distribution of MDA-MB-231 cells following treatment was determined (Figure 2D). When treated with 4nM Docetaxel, the concentration yielded maximum apoptosis, analysis of DNA content 48 hr following treatment revealed a significant proportion of cells with DNA content that diverged from the 2N peak, indicating extensive loss or gain of DNA content during aberrant cell division. Cells with severe chromosome loss are unviable (Kops et al., 2004), which explains the massive apoptosis induced by lower concentrations (4, 16nM) of Docetaxel. By contrast, these cells were not observed in cells treated with higher concentration of Docetaxel (64, 256nM) which mainly result in tetraploid cells with 4N DNA content, indicating cytokinesis is blocked when cells were treated with higher concentration thus the lethal effect of chromosome loss is avoided. A small percentage of cell with 8N DNA content was also observed, indicating endoreplication.

Several fates have been described for cells that undergone mitotic slippage, including apoptosis in interphase, cell-cycle arrest, and proliferate as polyploid cultures. Substantial evidence supports the theory that p53 restrains cell-cycle progression following exit from mitotic arrest (Gascoigne and Taylor, 2009; Lanni and Jacks, 1998). Thus MDA-MB-231 cells, which contain mutant p53 (Morse et al., 2005), may continue to proliferate and develop drug resistance.

To determine whether higher concentrations of Docetaxel are more likely to cause drug resistance in MDA-MB-231 cells, we detected the long-term cellular response by performing colony outgrowth assays. MDA-MB-231 were treated with increasing concentrations of Docetaxel for 36h (Figure 2E) or 48h (Figure S2B), and then cells were cultured in drug-free medium to allow for colony outgrowth. We found treatment of higher concentration did not yield more colonies. On the contrary, the number of colonies was negatively correlated with drug concentration, consistent with HeLa cells. Thus, cell killing assessed in short term assays does not predict the long-term cellular response to Docetaxel.

### Overexpression of Bcl-2/Bcl-xL suppresses Docetaxel induced apoptosis but does not promote clonogenic survival of tumor cells

Expression of antiapoptotic proteins was frequently found to be involved in the chemo-resistant phenotypes of human tumors cells. Moreover, following prolonged mitotic arrest induced by taxanes, the antiapoptotic functions of Bcl-2/Bcl-xL were reported to be abrogated by mitotic phosphorylation, thus coupling mitotic arrest to apoptosis (Terrano et al., 2010). Accordingly, expression of phosphor-mimetic Bcl-xL was unable to block mitotic death (Eichhorn et al., 2013). On the other hand, the antiapoptotic function of Surivivn was depend on mitotic phosphorylation which increase its stability (O’Connor et al., 2002), and a phosphorylation-defective Survivin T34A mutant was reported to exhibit a dominant-negative effect by initiating massive mitochondrial-dependent apoptosis in a variety of tumor cell lines (Mesri et al., 2001).

To verify the role of these apoptotic regulators in cellular response to Docetaxel, we first detected their endogenous expression (Figure 3A) and phosphorylation (Figure 3B) during mitotic arrest induced by Docetaxel. Cell-cycle-dependent expression of Survivin has been reported by many groups, which peaks at mitosis, correlated with its role in mitotic regulation. In accordance with previous observations (O’Connor et al., 2002), elevated expression of Survivin was observed in mitotic arrested cells, but not in cells that had exited mitosis (MDA-MB-231 cells at 24 h), though the massive concomitant apoptosis in HeLa cells suggest it’s not cytoprotective (Figure 1C). MDA-MB-231 cells did not express Bcl-2 but expressed higher level of Bcl-xL than HeLa cells. In accordance with the literature, phosphorylation of these apoptotic inhibitors was observed in cells arrested in mitosis (HeLa cells at 16, 24 h; MDA-MB-231 cell at 16 h), but not in cells before mitotic entry or that had exited mitosis (MDA-MB-231 cells at 24 h).

**Figure 3.**
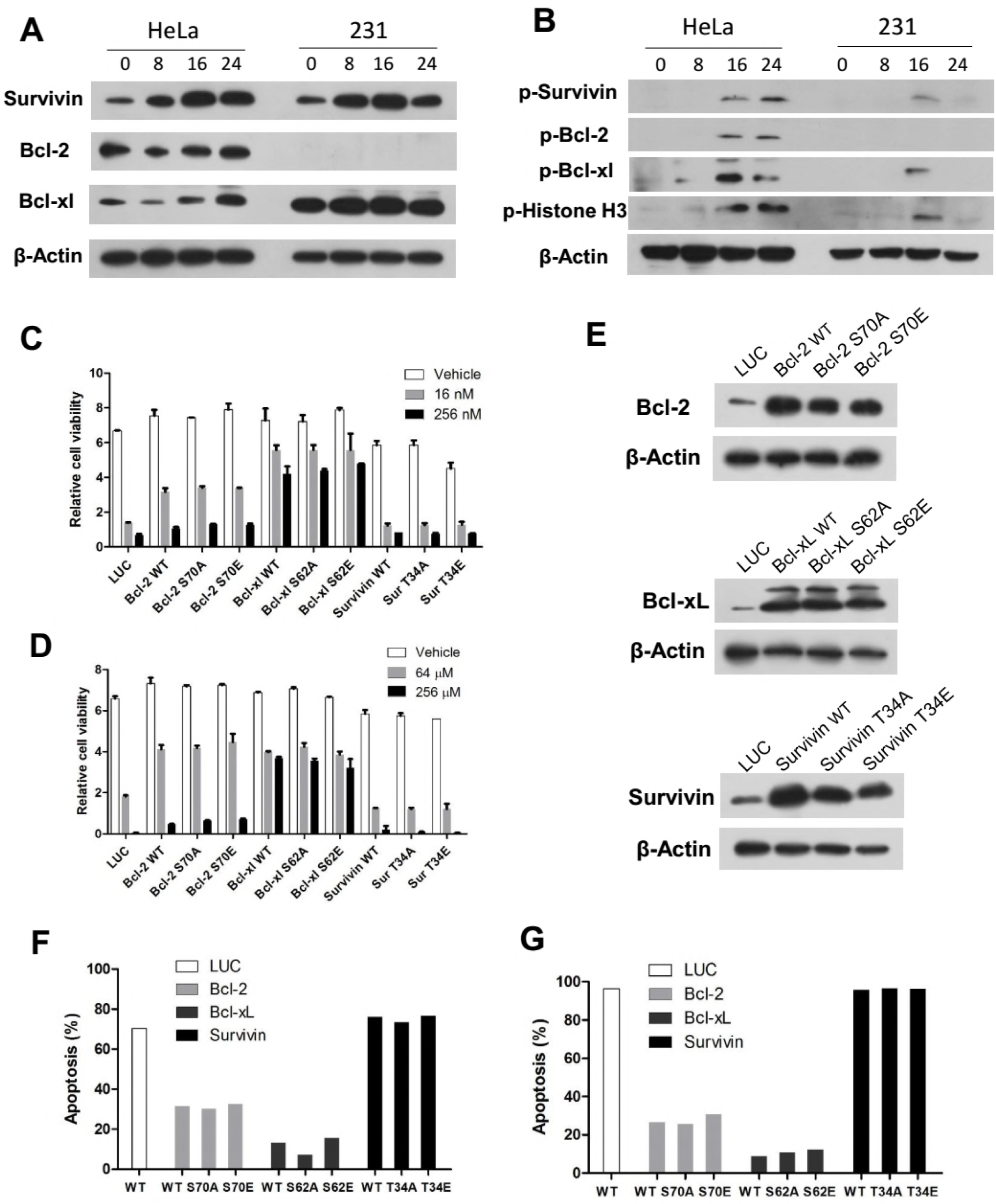
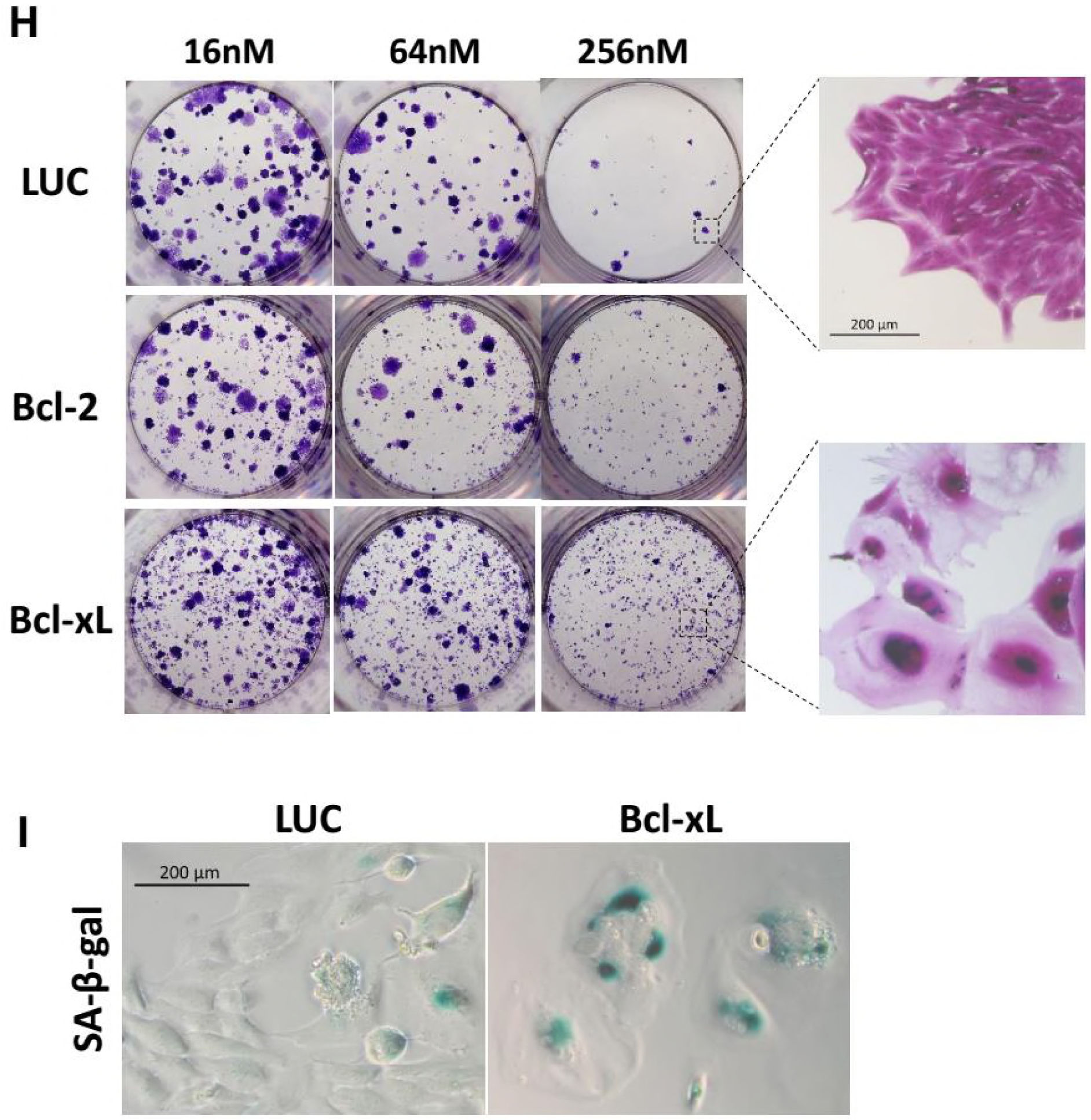
Overexpression of Bcl-2/Bcl-xL suppresses Docetaxel induced apoptosis but does not promote clonogenic survival. (A and B) Synchronized HeLa and MDA-MB-231 cells were treated with docetaxel. Cells were harvested at the indicated time intervals and analyzed for expression (A) and phosphorylation (B) of Survivin, Bcl-2, and Bcl-xl by Western blotting. Phosphorylation of histone H3 was used as a marker of mitosis. (C and D) HeLa cells were infected with lentiviral vectors expressing cDNA encoding Bcl-2, Bcl-xL, Survivin, or their phosphomimetic or deficient mutants for 48 h, followed by docetaxel (C) or cisplatin (D) treatment for another 48 h before cell viability was determined. Values represent the means ± S.D. (n = 2 wells). Lentiviral vectors expressing cDNA encoding luciferase (LUC) was used as a control. (E) Overexpression of indicated protein was validated in stable cell lines which were generated by lentiviral vectors infection followed by selection with puromycin. (F and G) Stable cell lines overexpressing Bcl-2, Bcl-xL, Survivin or their mutants were treated with docetaxel (256 nM) (F, see Figure S3C for details) or cisplatin (256 μM) (G, see Figure S3D for details) for 36 hr, then collected and analyzed for induction of apoptosis. (H) Stable cell lines were treated with the indicated concentrations of docetaxel for 36h, and then cultured in drug-free medium for 2 weeks. Colony outgrowth was photographed following fixation and staining with crystal violet. Magnified images shows proliferative cells that can form a colony versus senescent cells that cannot. (I) Increase of cellular senescence in Bcl-xL-expressing cells followed by Docetxel treatment compared with control cells. Cells were treated as described in (H) and assayed for SA-β-Gal activity.

To determine whether over-expression of antiapoptotic proteins result in drug resistance, we took advantage of a lentiviral expression system which circumvent the cytotoxicity mediated by plasmid transfection and shows premium gene transfer efficiency. HeLa cells were incubated with lentiviral particles expressing Bcl-2, Bcl-xL, Survivin, or phosphomimetic or deficient mutants for 48 h, followed by treatment with Docetaxel (Figure 3C) or cisplatin (Figure 3D) for another 48 h before cell viability was detected. Consistent with previous observations, over-expression of wild-type (WT) or mutant forms of Bcl-2 or Bcl-xL enhanced cell survival following Docetaxel treatment (Figure 3C). Only Bcl-xL, but not Bcl-2, was able to inhibit the cell death induced by higher concentrations of docetaxel (256 nM), supporting its more potent antiapoptotic function (Simonian et al., 1997). Expression of phosphomimetic or deficient proteins yielded results similar to those of WT proteins. Notably, neither over-expression of WT, phosphomimetic, or deficient mutant form of Survivin were able to inhibit cell death induced by docetaxel or cisplatin. Similar results were obtained with MDA-MB-231 cells (Figure S3A).

To validate these results, stable HeLa cell lines expressing WT or mutant forms of Bcl-2, Bcl-xL, or Survivin were generated by puromycin selection after lentivirus infection. Over-expression of the indicated protein was validated by western blotting (Figure 3E). Expression of the phosphomimetic or deficient mutant was validated by pyrosequencing of the cDNA for the indicated cell lines (Figure S3B). The induction of apoptosis was detected following docetaxel (Figure 3F and S3C) or cisplatin (Figure 3G and S3D) treatment. Similar to the cell viability assay, Bcl-xL was the most competent cell death inhibitor, and the proportion of apoptotic cells was also significantly reduced by Bcl-2. By contrast, over-expression of WT Survivin or its phosphomimetic or deficient mutants showed no effect on cellular sensitivity to either docetaxel or cisplatin.

A replication-deficient adenovirus encoding a non-phosphorylatable T34A mutant of Survivin was reported to induce spontaneous apoptosis and tumor cells sensitization to paclitaxel (Mesri et al., 2001). However, we found lentiviral expression of Survivin T34A mutant did not yield marked cell death or sensitization to docetaxel in HeLa cells (data not shown) as well as in MDA-MB-231 or MCF-7 cells (Figure S3E and S3F). The apparent contradiction with previous reports may be due to the lentiviral vectors used in this study which cause little or no disruption of the target cell even at high titer. Whereas Cellular toxicity following plasmid transfection or adenovirus infection at a high multiplicity of infection (MOI) has been reported (Brand et al., 1999).

While apoptosis was clearly blocked by overexpression of Bcl-2/Bcl-xL, colony outgrowth assays were carried out to determine the long-term effect of the expression antiapoptotic proteins. HeLa cells stably expressing Bcl-2, Bcl-xL, or LUC control were treated with Docetaxel for 36h and then allowed to grow in drug-free medium for 2 weeks. We found overexpression of Bcl-2/Bcl-xL did not lead to increased clonogenic survival (Figure 3H), consistent with a previous report (Yin and Schimke, 1995). The number of colonies correlated with drug concentration rather than the expression of antiapoptotic proteins. At the same concentration, expression of Bcl-2/Bcl-xL did not yield more colonies than control cells, but it did yield more viable cells that lose the ability to proliferate as they failed to grow into colonies (Figure 3H right). These cells adopted flattened and enlarged morphology characteristic of cellular senescence, which was confirmed by senescence-associated β-galactosidase (SA-β-Gal) staining (Figure 3I). Since cellular senescence irreversibly arrests cell division, it is considered an important form of cell “death” which contribute to the deletion of cancer cells (Chang et al., 1999). The switch of apoptosis to senescence may explain why overexpression of Bcl-2/Bcl-xL protected cell viability against therapy but failed to promote cell growth after drug withdraw.

### Depletion of Survivin suppresses apoptosis through facilitating mitotic slippage by compromising the SAC

Increased expression of Survivin in cancer has been reported by numerous groups, which is correlated with taxanes resistance both in vitro (Li et al., 1998) and in vivo (Zaffaroni et al., 2002). Since overexpression of Survivin failed to inhibit apoptosis, to verify the antiapoptotic role of Survivin, we depleted its expression by using RNA interference (RNAi). Depletion of Survivin was reported to result in spontaneous apoptosis as well as sensitization to chemotherapeutic agents (Zaffaroni et al., 2005). By using different siRNA sequence designed by biotech companies or from previous literature, we found the proportion of apoptosis following transfection of individual siRNAs varied widely. While some siRNAs induced substantial apoptosis, the level of apoptosis induced by others was only slightly more than that of control siRNA (Figure S4A). The cell death was found to occur following aberrant mitosis as indicated by accumulation of multinucleated cells (data not shown), correlating with the crucial role of Survivin in mitotic regulation (Mita et al., 2008). Moreover, we found 2’-O-methyl modification of residues in siRNA strand (Jackson et al., 2006) significantly reduced toxicity of siRNA against Survivin, while gene silencing was not markedly affected (Figure S4B and S4C). The fraction of apoptosis following siRNA transfection was also found to be reduced by single base substitution (Figure S4D and S4E), suggesting dell death was due to nonspecific toxic effects. The siRNA sequence that exhibit minimal toxicity (Ana et al., 2003) was used for following experiments.

To detect whether depletion of Survivin sensitize tumor cells to Docetaxel, HeLa cells were incubated with siRNA against Bcl-xL or Survivin, induction of apoptosis and cell viability were determined following Docetaxel treatment. Transfection of siRNA against Bcl-xL clearly sensitized HeLa cells to Docetaxel as shown by elevated level of apoptosis (Figure 4A and S4F) and diminished level of cell viability (Figure 4B) following treatment with different concentrations of Docetaxel. Unexpectedly, depletion of Survivin suppressed apoptosis induced by Docetaxel, which was more pronounced when cells were treated with 16nm rather than 64nM or 256nM Docetaxel. Note that depletion of Survivin also led to enhanced viability of MCF-7 cells following Docetaxel treatment (Figure S4G and S4H).

**Figure 4.**
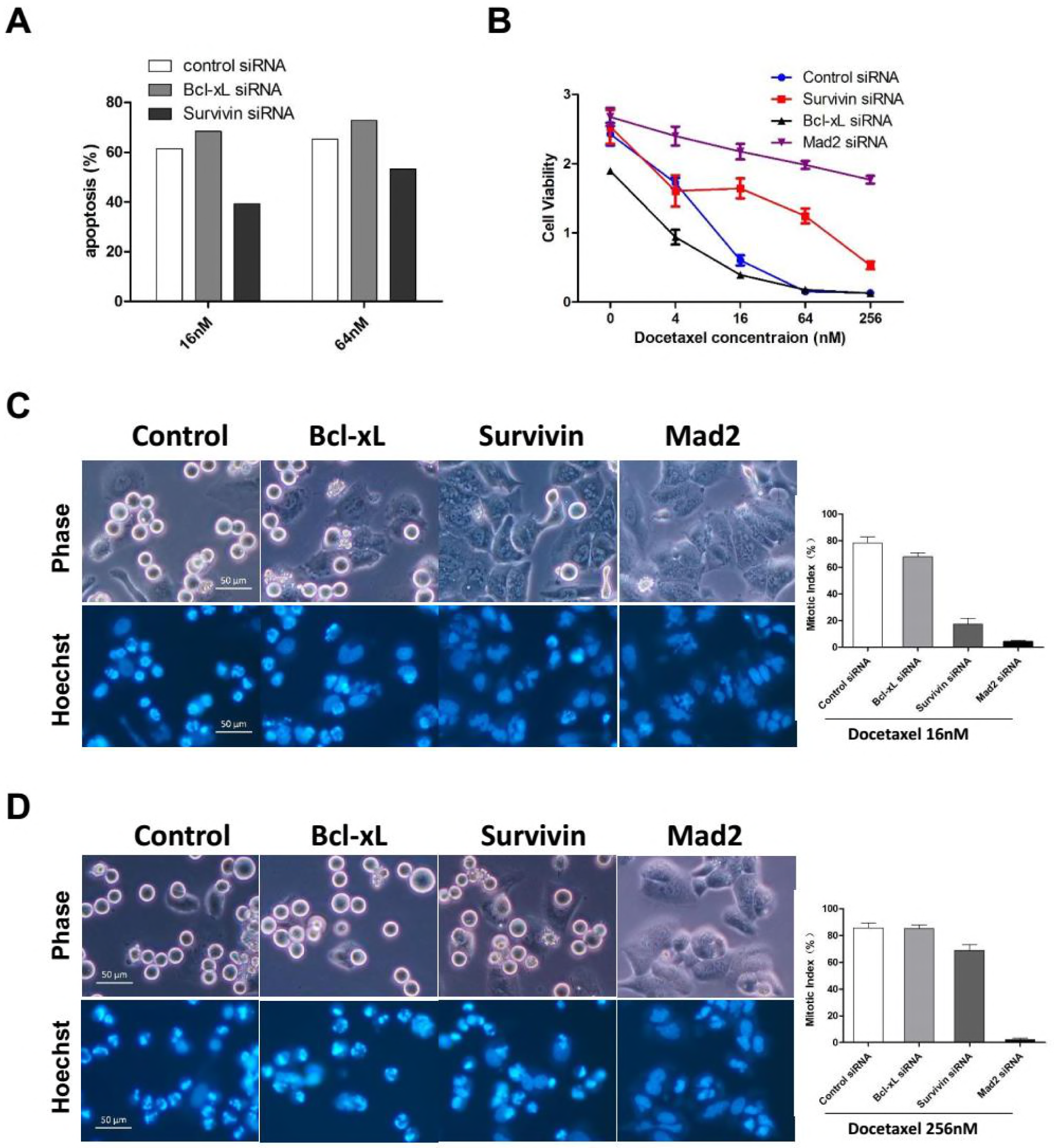
Depletion of Survivin suppresses apoptosis induced by Docetaxel. (A) HeLa cells were incubated with the indicated siRNAs for 36 hr followed by 16nM or 64nM Docetaxel treatment for 24h. Cells were collected and analyzed for induction of apoptosis (see Figure S4F for details). (B) HeLa cells were incubated with the indicated siRNAs for 36 hr followed by treatment with increasing concentrations of Docetaxel for 48 hr before cell viability was determined. Values represent the means ± S.D. (n = 3 wells). (C and D) HeLa cells were incubated with the indicated siRNAs for 36 hr and synchronized by thymidine block. Following a 16 hr exposure to 16nM (C) or 256nM (D) Docetaxel, cells were stained with Hoechst 33342 and photographed using phase and Hoechst fluorescence. Mitotic indexes were determined by counting the percentage of cells with spherical morphology and condensed chromatin. Values represent the means ± S.D. (n = 3 fields).

We have shown the rapid death of HeLa cells following prolonged mitotic arrest, which depend on the chronic activation of SAC. As an essential regulatory subunit of the chromosome passenger complex (CPC), Survivin was shown to be required for SAC function in the presence of taxane (Ana et al., 2003). To test whether Survivin-depleted cells arrested normally at mitosis following Docetaxel treatment, synchronized cells were exposed to 16nM (Figure 4C) or 256nM (Figure 4D) Docetaxel for 16 hr, and then mitotic cells or cells undergone mitotic slippage were observed and photographed under a fluorescence microscopy and mitotic index was calculated. While majority of cells transfected with control or Bcl-xL siRNA arrested normally, cells tranfected with siRNA against Mad2, the central components of the SAC, failed to block in mitosis following exposure to either 16nM or 256nM Docetaxel, indicating abolishment of SAC through depletion of Mad2.

The depletion of Survivin has a more complicated effect on the mitotic arrest following Docetaxel treatment. Although the mitotic index was only slightly reduced in Survivin-depleted cells when treated with 256nM Docetaxel. The majority of Survivin-depleted cells failed to block in mitosis by treatment of 16nM Docetaxel, contrasting with control or Bcl-xL-depleted cells, indicating a compromised SAC response due to depletion of Survivin. Similar observations have been made by a previous study (Z et al., 2008). Thus, Survivin-depleted cells are more inclined to undergo mitotic slippage following Docetaxel treatment, through which they escape the death due to prolonged mitotic arrest. Taken together, we have not observed any anti-apoptotic effects of Survivin following Docetaxel treatment.

### Inhibition of the SAC suppresses apoptosis but enhances the long-term efficacy of different microtubule toxins

Although inhibition of SAC by depletion of Survivn or Mad2 led to resistance to Docetaxel induced apoptosis, we have demonstrated that short-term cell death was not necessarily associated with long-term cellular responses after drug withdraw. To determine whether SAC inhibition lead to Docetaxel resistance, colony formation assay was performed which was more reflective of therapeutic response in vivo (Brown and Attardi, 2005). Cultures transfected with siRNA against Survivin, Mad2 or Aurora B, which is the central kinase of the CPC, were treated with Docetaxel, and then allowed to grow in drug-free medium for 2 weeks before fixed and stained for colony outgrowth.

Surprisingly, suppression of Survivin, Aurora B or Mad2 did not lead to drug resistance. On the contrary, the colony outgrowth of tumor cells was almost completely blocked by depletion of Survivin or Mad2 (Figure 5A upper panels). Depletion of Aurora B shows a lesser effect on suppression of colony formation, possibly due to the inferior knockdown efficiency of the siRNA targeting Aurora B compared with that targeting Survivin or Mad2 (Figure S5A). The blockage of colony outgrowth was not completely attributed to cell death, we found many Survivin or Mad2 depleted cells remained alive at the time of fixation but lose the ability to proliferate with morphological characteristics of cellular senescence which was confirmed by increased SA-β-Gal activity (Figure 5A lower panels). Notably, overexpression of antiapoptotic proteins was also found to increase cellular senescence following Docetaxel treatment, however, colony outgrowth of tumor cells was blocked by depletion of Survivin/Mad2 rather than overexpression of Bcl-2/Bcl-xL, indicating cellular senescence alone was not sufficient to inhibit long-term growth of tumor cells.

**Figure 5.**
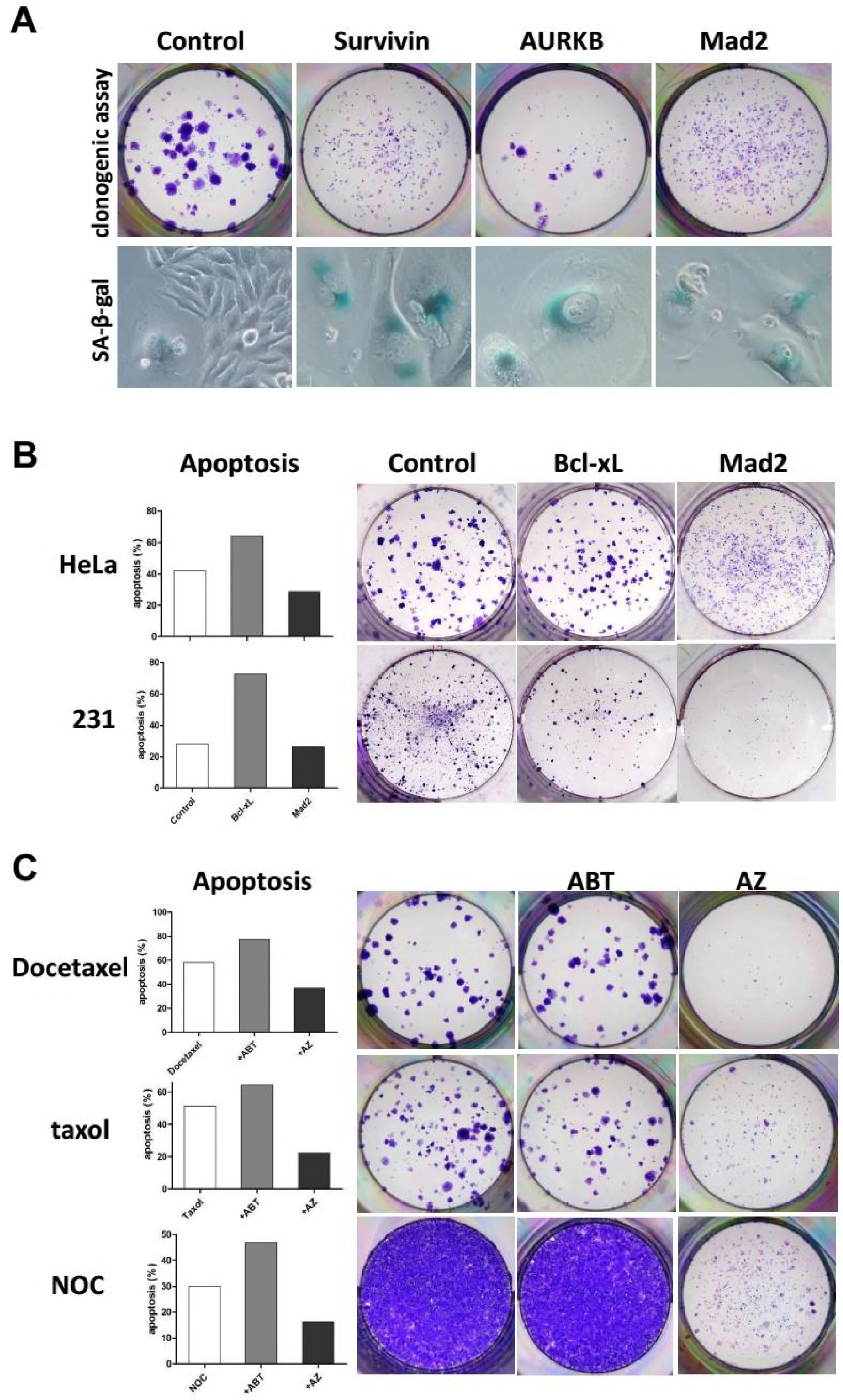
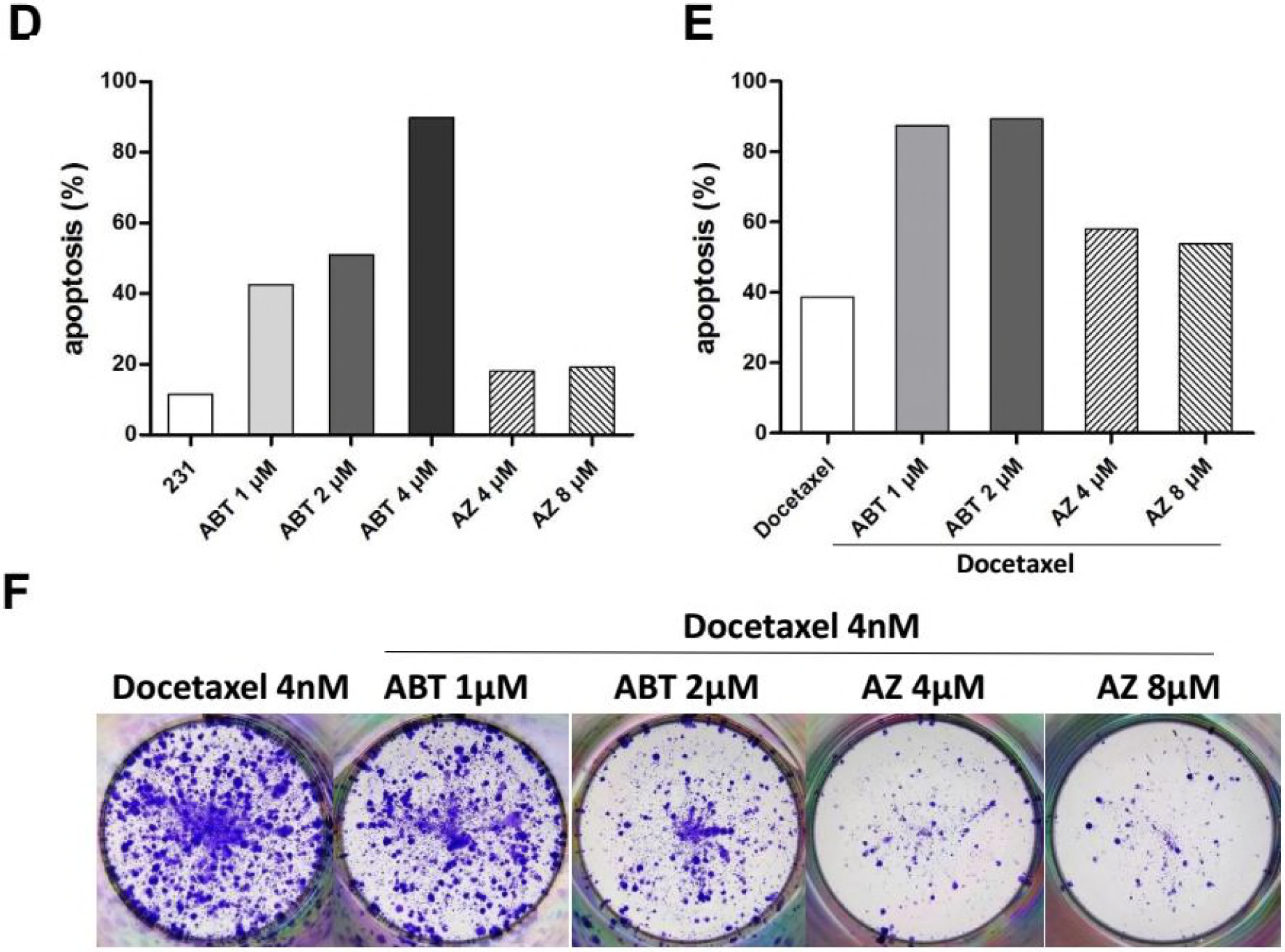
Inhibition of the SAC suppresses apoptosis induced by microtubule toxins but enhances their long-term efficacy. (A) HeLa cells were incubated with the indicated siRNAs for 48 hr followed by Docetaxel treatment for 36h, and then cultured in drug-free medium for 2 weeks. Cells were either stained and photographed for Colony outgrowth (upper panels) or assayed for senescence (lower panels). (B) HeLa and MDA-MB-231 cells were incubated with indicated siRNAs for 48h and then exposed to Docetaxel for 36h, cells were either collected immediately and analyzed for induction of apoptosis (left, see Figure S5B for details), or cultured in drug-free medium to allowed for Colony outgrowth (right). (C) HeLa cells were treated with microtubule toxins, including Docetaxel, paclitaxel or nocodazole, alone or in combination with ABT-737 or AZ3146, and then either collected and analyzed for induction of apoptosis (left, see Figure S5C for details), or cultured in drug-free medium to allow for Colony outgrowth (right). (D) MDA-MB-231 cells were treated with the indicated concentrations of ABT-737 or AZ3146 for 36h, then collected and analyzed for induction of apoptosis (see Figure S5G for details) (E and F) MDA-MB-231 cells were first treated with the indicated concentrations of ABT-737 or AZ3146 for 24h, and then with ABT-737 or AZ3146 in combination with Docetaxel (4nM) for 36h. Cells were either collected and analyzed for induction of apoptosis (E, see Figure S5K for details), or cultured in drug-free medium to allow for colony outgrowth (F).

The antiapoptotic Bcl-2-family proteins have been regarded as potential drug targets, inhibition of these proteins was reported to synergize with taxanes, leading to tumor regression in xenographs (Ealovega et al., 1996) and improved survival of patients (Webb et al., 1997). Here we compared the effect of targeting antiapoptotic Bcl-xL or SAC component Mad2 on cellular response to Docetaxel. HeLa and MDA-MB-231 cells were incubated with siRNA against Bcl-xL or Mad2 for 48 hr followed by exposure in Docetaxel for another 48 hr, and then either harvested immediately for detection of apoptosis (Figure 5B left) or allowed to grow in drug free medium for outgrowth of colony (Figure 5B right). Sensitization to Docetaxel was achieved by depletion of Bcl-xL rather than Mad2 in both HeLa and MDA-MB-231 cells, as judged by the elevated level of apoptosis following treatment, which was more pronounced in MDA-MB-231 cells (Figure 5B left and S5B). Whereas, the level of apoptosis was reduced by depletion of Mad2 in HeLa cells but not in MDA-MB-231 cells which cannot sustain a prolonged mitotic arrest. Surprisingly, clonogenic survival of tumor cells was nearly completely blocked by depletion of Mad2 rather than Bcl-xL in both Hela and MDA-MB-231 cells (Figure 5B right).

Since BH3-mimetics were reported to synergize with taxanes, we compared the antitumor effect of ABT-737 (Oltersdorf et al., 2005), a BH3 mimetic which bound with high affinity to Bcl-2, Bcl-xL, and Bcl-w, with AZ3146 (Hewitt et al., 2010), an inhibitor of MPS1 which is an essential SAC kinase, when used in combination with Docetaxel. HeLa cells were treated with 4µM ABT-737 or AZ3146 in combination with different microtubule toxins including Docetaxel, paclitaxel and nocodazole, and then either harvested by 48 hr for detection of apoptosis or allowed to form colonies in drug free medium. Similar to Bcl-xL depletion, ABT-737 sensitized Hela cells to different microtubule toxins (Figure 5C left and S5C) but the combined treatment failed to prevent colony formation, which was nearly completely blocked by the combination of AZ3146 and different concentration of microtubule toxins (Figure 5C right S5D S5E and S5F). It should be noted that prolonged exposure to paclitaxel (48 hr) or nocodazole (60 hr) in combination with AZ3146 was needed to block colony formation compared with Docetaxel (36 hr). At higher concentrations (64nM or 256nM), Docetaxel alone was sufficient to block colony formation (Figure S5D). That explains why the enhanced the efficacy of Docetaxel by AZ3146 was more pronounced at lower concentrations.

Compared with HeLa cells, MDA-MB-231 cells were markedly more sensitive to ABT-737 induced cell death. 4µM ABT-737 alone result in nearly 90 percent cell killing by 36 hr drug treatment (Figure 5D and S5G). Thus, the concentration of ABT-737 was reduced when used in combination with Docetaxel in MDA-MB-231 cells. At 1 or 2µM, 36h exposure to ABT-737 in combination with Docetaxel lead to substantial apoptosis (83% at 1µM ABT-737, 95% at 2µM ABT-737), compared with the cell death induced by Docetaxel alone (39%) (Figure S5H and S5I). Unexpectedly, despite the elevated level of apoptosis by ABT-737, the number of colonies was not markedly reduced following combination treatment compared with Docetaxel alone (Figure S5J). The combination of AZ3146 and Docetaxel, which led to moderate level of apoptosis (60% at 4µM AZ3146, 59% at 8µM AZ3146), was failed to block colony formation as well (Figure S5J).

Given that MDA-MB-231 cells have a slower doubling time compared with that of HeLa cells, suggesting an insufficient number of cells pass through mitosis during 36 hr treatment. We therefore extended the time that cells are exposed to AZ3146, as well as ABT-737. MDA-MB-231 cells were sequentially treated with AZ3146 or ABT-737 for 24 hr followed by 36 hr combination treatment with 4nM (Figure 5F) or 8nM (Figure S5L) Docetaxel, and then cells were allowed to grow in drug free medium. Under these conditions, AZ3146 showed a clear advantage over ABT-737 in suppression of colony formation when combined with Docetaxel (Figure 5F and S5L). Note that the sequential treatment with 2µM ABT-737 and Docetaxel yielded a lower level of apoptosis (89%) (Figure 5E and S5K) compared with single treatment for 36 hr (95%) (Figure S5H and S5I), that is because the cell death following the first 24 hr ABT-737 treatment was not counted. The true rate of apoptosis following sequential treatment with 2µM ABT-737 must be higher than 95%, a level much higher than apoptosis induced by AZ3146 in combination with Docetaxel (58% at 4µM AZ3146, 53% at 8µM AZ3146) or Docetaxel alone (38%) (Figure 5E and S5K). In addition, an increased level of cellular senescence was also observed following combination treatment with AZ3146 and Docetaxel (Figure S5M), although not as obvious as Mad2-depleted cells.

Together, inhibition of SAC exerts superior efficacy against clonogenic survival of both SAC proficient HeLa cells and less proficient MDA-MB-231 cells to apoptosis sensitizer, when combined with different microtubule toxins.

### Partial abrogation of SAC synergizes with Docetaxel to block colony formation

SAC components were suggested to play different roles in cellular response to taxane. While partial depletion of Mps1 or BubR1 sensitizes tumor cells to paclitaxel, reduced levels of Mad2 failed to do so (Janssen et al., 2009), instead, it was associated with drug resistance (Sudo et al., 2004). Depletion of Mad2 was reported to not only block cell death, but also lead to enhanced colony outgrowth following paclitaxel treatment (Bargiela-Iparraguirre et al., 2014; Prencipe et al., 2009), which runs directly counter to our observation. To clarify the effect of Mad2 depletion, we used another siRNA sequence from a previous literature (Kabeche and Compton, 2012) to target Mad2 (siRNA #2). The aforementioned siRNA against Mad2 used in this study was referred to as siRNA #1. Transfection of either siRNA led to abrogation of mitotic arrest (Figure 6C) and induction of cellular senescence (Figure S6A) when combined with Docetaxel. However, clonogenic survival of HeLa cells following Docetaxel treatment was only blocked by siRNA #1 but not by siRNA #2 (Figure 6A).

**Figure 6.**
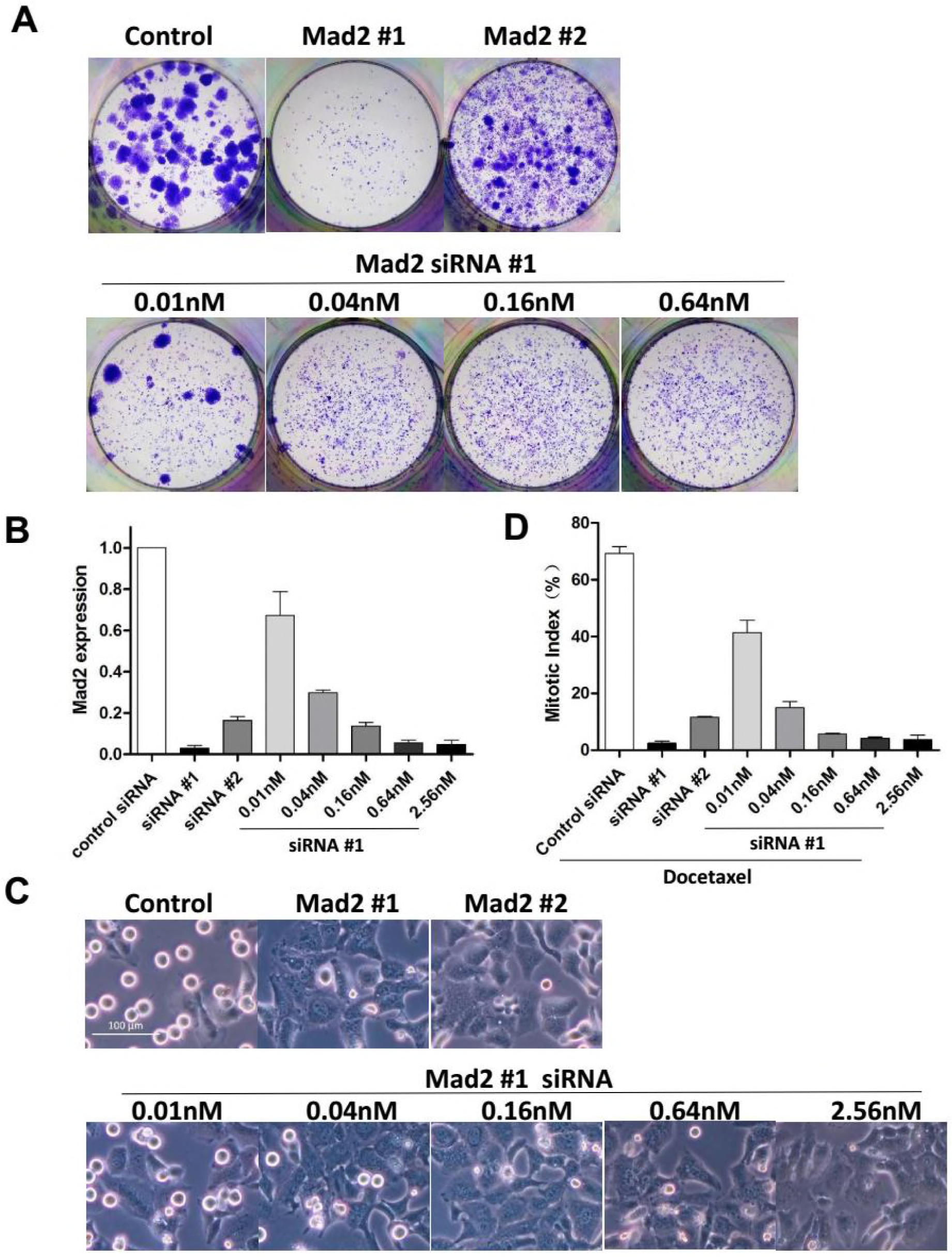
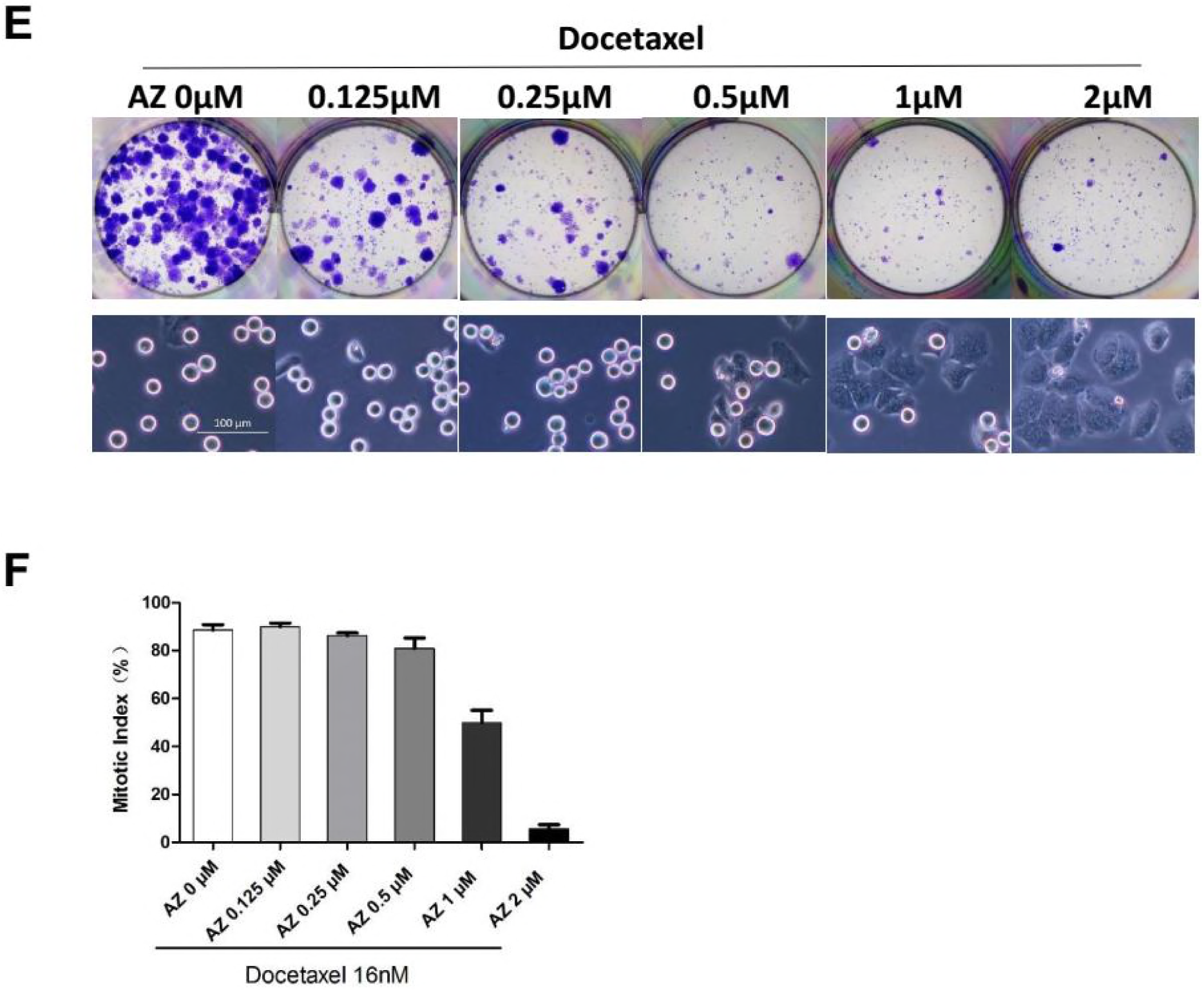
Partial Abrogation of SAC synergizes with Docetaxel in blocking colony formation. (A) HeLa cells were transfected with the indicated concentrations of siRNA for 48h followed by Docetaxel treatment for 36h, and then cultured in drug-free medium for 2 weeks. Colony outgrowth was photographed following fixation and staining with crystal violet. (B) HeLa cells were incubated with the indicated concentrations of siRNA for 48hr, then harvested and subjected to quantitative real-time PCR analysis for Mad2 expression. Values represent mean ± s.d. (n = 2 independent experiments). (C and D) Cells were incubated with the indicated concentrations of siRNAs for 48 hr and synchronized by thymidine block. Following a 16 hr exposure to Docetaxel the cells were photographed (C) and mitotic index (D) was determined by counting the percentage of cells with spherical morphology. Values represent the means ± S.D. (n = 3 fields). (E and F) HeLa cells were treated with the indicated concentrations of AZ3146 in combination with Docetaxel. Following treatment the cells were either photographed (E, lower panels) to determine mitotic index (F), or cultured in drug-free medium to allow for Colony outgrowth (E, upper panels).

Note that the knockdown efficiency of siRNA #2 was inferior to siRNA #1(Figure 6B), which could mean a partial depletion of Mad2 through transfection of siRNA #2, contrasting with complete depletion by siRNA #1. While complete loss of Mad2 alone was reported to result in cell lethality (Michel et al., 2004), partial depletion of Mad2 which results in a defective SAC is compatible with life and is associated with resistance to paclitaxel (Sudo et al., 2004). To determine the influence of the level of Mad2 to cellular response to Docetaxel, we reduced the level of MAD2 transcripts to different degrees by transfection of serial diluted siRNA #1 (Figure 6B). We found the mitotic index at 16h following Docetaxel treatment (Figure 6D) was correlated with the level of Mad2 transcripts (Figure 6B), consistent with the critical role of Mad2 in mitotic checkpoint control. At a concentration of 0.04nM, transfection of siRNA #1, which lead to 70% reduction of Mad2 transcript, dramatically reduced colony formation when combined with Docetaxel (Figure 6A). Whereas, transfection of RNA #2 (40nM) which lead to 80% reduction of Mad2 transcript, failed to block colony formation, suggesting that the failure to block colony formation by siRNA #2 was unrelated to the level of Mad2 transcripts.

To exclude the possibility that suppression of colony formation by combination treatment was due to Mad2 depletion alone, we determined colony outgrowth following transfection of siRNA #1 without Docetaxel treatment. We found colony outgrowth can be blocked by transfection of higher concentration of siRNA (≥0.64nM) which lead to complete depletion of Mad2 (Figure S6B). Transfection of lower concentration (≤0.16nM) failed to completely block colony formation (Figure S6B) unless in combination with Docetaxel (Figure 6A), suggesting a synergistic effect between Docetaxel and partial abrogation of SAC.

To further address this, increasing concentrations of MPS1 inhibitor AZ3146 were used in combination with Docetaxel. At concentrations ≥1μM, mitotic index following Docetaxel treatment was markedly reduced by AZ3146 (Figure 6F) which facilitate mitotic slippage without cell division, resulting multinucleated cells in the subsequent interphase (Figure 6E lower panels). At concentrations ≤0.5μM, mitotic arrest was barely affected by AZ3146, suggesting the spindle checkpoint response was not severely compromised. Notably, when combined with Docetaxel, even the lowest concentration (0.125μM) of AZ3146 led to marked reduction of colony formation compared with Docetaxel treatment alone (Figure 6E upper panels), indicating that even partial abrogation of SAC could synergize with docetaxel to block the long-term growth of tumor cells. Note that exposure to AZ3146 alone (up to 8μM, for 48 hr) was not sufficient to block colony formation (Figure S6C).

## Discussion

### Mitotic arrest following taxane treatment and determinant of cell fate

Taxanes have been successfully used in the clinic for years, but their mechanism of antitumor action remains unclear. Since substantial cell death is often observed following prolonged mitotic block, it is easy to attribute its antitumor effects to mitotic arrest, with cells that arrest longer being more likely to die. In line with this notion, the accumulation of proapoptotic signals during prolonged mitotic arrest has been observed by many studies (Gascoigne and Taylor, 2008) and blocking mitotic exit have been proposed as a powerful strategy to eliminate tumor cells by allowing more time for death initiation (Huang et al., 2009). Conversely, Mitotic slippage is thought to be one of the main mechanisms by which cancer cells avoid mitotic death and develop drug resistance (Manchado et al., 2012).

Consistent with previous studies, we find the fraction of apoptotic cells increases as the cells undergone prolonged mitotic arrest, and promotion of mitotic slippage is cellular protective which markedly reduced the cell death following 24 hr Docetaxel treatment. On the other hand, similar levels of apoptosis were observed at 50 hr after Docetaxel treatment between cells undergone prolonged mitotic arrest and mitotic slippage, suggesting promotion of mitotic slippage only delays cell death induced by Docetaxel, by switching apoptosis from mitosis to the subsequent interphase.

Moreover, promotion of Mitotic slippage did not enhance the long-term cell growth, as detected by clonogenic assay which is more relevant to tumor response in vivo. These findings indicate that although the prolonged mitotic arrest is associated with apoptotic cell death, facilitating mitotic exit does not promote Docetaxel resistance.

In addition, we found treatment with higher concentration of Docetaxel always yields better inhibition of colony outgrowth, even in cells undergo mitotic slippage, suggesting the superior efficacy of higher concentration of Docetaxel cannot be ascribed to prolonged mitotic arrest. Since Taxanes target and stabilize microtubules, thereby preventing normal spindle assembly and chromosome segregation. It’s reasonable to infer that Docetaxel of higher dose may lead to more severe chromosome segregation errors compared with lower concentrations. We therefore hypothesize that if a cell does not undergo death during drug induced mitotic arrest, its fate may depend on the severity of chromosome segregation errors, and each cell may contain a threshold for the amount of errors it can stand, beyond which the cell is destined to die, sooner or later. In line with this notion, it was recently reported that clinically relevant concentrations of paclitaxel cause cell death due to chromosome mis-segregation on multipolar spindles rather than mitotic arrest (Zasadil et al., 2014), and elevating the frequency of chromosome mis-segregation has been proved to be an effective strategy to sensitize tumor cells to taxane (Janssen et al., 2009).

### Mode of cell death following therapy: Apoptosis versus senescence

Although multiple forms of cell death following therapy has been reported, it is generally believed that conventional cytotoxic agents including taxanes trigger the cell death via activation of the intrinsic apoptosis, a pathway involving mitochondrial outer membrane permeabilization (MOMP). Failure to initiate apoptosis appears to be an important mechanism through which tumor cells develop drug resistance (Adams and Cory, 2007).

Due to the short time scale of apoptosis, the amount of cell death or cell viability is often assessed within 48 hr following treatment (Sudo et al., 2004; Swanton et al., 2007). However, delayed cell death is common following therapy. Especially when the clinical relevant lower concentrations of paclitaxel is used, which lead to chromosome misdistribution rather than sustained mitotic arrest (Zasadil et al., 2014). Contrast to the rapid cell death following mitotic arrest, the amount of segregation errors accumulates as cells undergo subsequent mitoses, which ultimately resulted in lethality within 2–6 cell divisions (Kops et al., 2004). Thus, the cell death could be severely underestimated if assayed at a fixed time point. Accordingly, overexpression of antiapoptotic proteins was found by us and others to only delays drug-induced apoptosis but does not increase clonogenic survival after Drug Treatment (Yin and Schimke, 1995).

Furthermore, apoptosis assay measures the rate rather than the overall level of cell killing (Brown and Attardi, 2005). Whereas, due to the heterogeneous nature of tumor, it might include a diverse collection of cells with differential levels of sensitivity to treatment (Dagogo-Jack and Shaw, 2018). The treatment response was not always determined by the majority of cells that are sensitive to therapy, because only a small fraction of cells that survive and retain the ability to proliferate could repopulate the tumor after drug withdraw (Kim and Tannock, 2005). By using clonogenic assay which measures overall level of cell death as well as the ability to proliferate, we found the proportion of cell death detected by short-term assays does not predict the long-term cellular response. For example, despite the fact that ABT-737 increased the apoptosis of HeLa cells following Docetaxel treatment by 20%, this strategy failed to diminish the number of colony outgrowth compared with Docetaxel treatment alone. Similarly, when used in MDA-MB-231 cells, ABT-737 elevated the apoptosis rate following Docetaxel treatment from 39% to 95%, however, this combination of agents still failed to block colony outgrowth. These data indicate that the level of apoptosis following treatment can be misleading in assessing the long-term cellular response to taxanes.

Notably, apoptotic cell death was also suggested to promote cancer (Ichim and Tait, 2016). Although evasion of apoptosis is considered to be a hallmark of cancer, cancer cells are often not inherently apoptosis resistant. Apoptosis has been observed in a wide array of untreated patient tumors and contributed to the high rate of cell loss in malignant tumors (Lowe and Lin, 2000; Weaver, 2014). emerging evidence indicate that apoptosis could promote cancer through induction of compensatory cell proliferation of neighbouring cells (Huang et al., 2011). Several other mechanisms have also been implicated in this process (Ichim and Tait, 2016). Along this line, higher levels of apoptosis have also been shown to correlate with poorer prognosis in some cancer types. On the contrary, antiapoptotic proteins expression correlate with better prognosis in certain cancers.

In contrast to apoptosis, the presence of senescent cells was found exclusively in premalignant lesions but not in malignant ones (Collado et al., 2005), suggesting a role for senescence as a barrier to tumor progression (Collado and Serrano, 2010). Cellular senescence is an irreversible program of cell-cycle arrest that can be induced by diverse stimuli leading to the permanently loss of proliferative capacity of cells. Of note, tumor models with inactivated apoptotic signaling pathways respond to senescence-inducing drugs (Ewald et al., 2010). Thus, drugs aimed at selectively inducing cellular senescence could represent a promising novel approach for solid tumors that have diminished apoptotic potential.

The correlation between aberrant levels of SAC components and induction of cellular senescence has been observed by several groups (Bargiela-Iparraguirre et al., 2014; Lentini et al., 2012; Prencipe et al., 2009), and the senescent phenotype was further enhanced following paclitaxel treatment (Bargiela-Iparraguirre et al., 2014). Here we show that SAC inhibition switches Docetaxel-induced apoptosis to senescence, and the combined treatment of Docetaxel and SAC inhibition shows superior efficacy against clonogenic survival of tumor cells to Docetaxel treatment alone. However, the superior anti-tumor effect cannot be completely attributed to the induction of cellular senescence, for there are situations that enhanced cellular senescence is insufficient to block colony outgrowth. For example, overexpression of antiapoptotic Bcl-xL switched Docetaxel-induced apoptosis to senescence, whereas colony outgrowth after drug withdraw was barely affected. The elevated level of cellular senescence may reflect the enhanced pro-survival signals or the diminished cell death signals, in combination with the permanent loss of proliferative capacity of tumor cells. The senescence phenotype following SAC inhibition was probably a manifestation of severe chromosome distribution errors that beyond the threshold cells can withstand. Further investigations are needed to clarify the contribution of cellular senescence to treatment outcomes.

### False-positive phenotypes due to nonspecific activity of RNA interference

Consistent with the role of Survivin in apoptosis inhibition, depletion of Survivin has been reported to induce spontaneous apoptosis as well as sensitization to anticancer agents (Zaffaroni et al., 2005). However, by using multiple individual siRNAs targeting Survivin, we found substantial apoptosis was only observed following transfection of 2 out of 4 Survivin siRNAs, the other 2 siRNAs mainly led to mitotic abnormities as judged by accumulation of multinucleated cells, which may ultimately result in cell death rather than the rapid induction of apoptosis. Notably, the cell death induced by siRNA is sequence dependent, which could be reduced by base mismatch.

Furthermore, 2’-O-methyl modification (Jackson et al., 2006) significantly reduced the toxicity of Survivin siRNAs without compromising gene silencing, which strongly indicate that the observed cell death following siRNA transfection is due to non-specific toxic effect. Indeed, the widespread off-target gene modulation has been reported to induce a concentration dependent apoptotic phenotype (Fedorov, 2006), thus is considered to be a major hurdle to the use of RNAi.

In addition, when combined with Docetaxel, we found transfection of siRNA against Mad2 (#1) synergize with Docetaxel to block colony formation of tumor cells. By contrast, another siRNA (#2) targeting Mad2 failed to do so. Initially, we attribute this to the difference in gene silencing efficiency between siRNA #1 and #2. However, when the similar level of MAD2 transcripts was achieved, we found colony outgrowth was blocked by transfection of diluted siRNA #1 rather than undiluted siRNA #2, indicating nonspecific activity contribute to the failure of siRNA #2 to block colony formation.

Sensitization to paclitaxel was reported to be achieved by reducing levels of Mps1 or BubR1, but not by reducing Mad2 (Janssen et al., 2009). Suppression of Mad2 has been associated with paclitaxel resistance shown by enhanced clonogenic survival of tumor cells transfected with Mad2 siRNA (Bargiela-Iparraguirre et al., 2014; Prencipe et al., 2009), which was contrary to our observations. Our finding suggests that non-specific activity of siRNA may exist in a large number of studies and thus affect the reliability of their conclusions.

### Spindle assembly checkpoint: a survival checkpoint in cancer

SAC is a conserved surveillance mechanism by which cell prevent chromosome mis-segregation. A compromised SAC may lead to CIN and aneuploidy, both of which are common features of human cancers and are associated with intrinsic taxane resistance (Swanton et al., 2009). Although CIN cells was assumed to have a defective or weakened SAC, time-lapse imaging of multiple cancer and non-transformed cell lines revealed that CIN cells arrested for as long as—and even longer than non-transformed cells (Gascoigne and Taylor, 2008), indicating the SAC in CIN cells is clearly functional. Moreover, mutations in spindle checkpoint genes are rare in human tumors and the SAC components are not downregulated but rather overexpressed in most chromosomally unstable tumor cell lines and tissue samples (Yuan et al., 2006). For example, Mad2 overexpression is common in human tumors, and has been regarded as a critical mediator of the CIN phenotype (Schvartzman et al., 2011). Notably, Mad2 overexpression led to a hyperactive SAC which was suggested to play a causal role in cancer initiation and progression (Sotillo et al., 2007). This notion was clearly contradictory to the role of SAC as a safe guardian of genomes.

The observation that even a single unattached kinetochore can delay mitosis by many hours (Rieder et al., 1995), suggests that the SAC generates a switch-like ‘all-or-none’ response. However, emerging evidence shows that SAC signal is graded rather than binary, and its strength correlates with the number of unattached chromosomes (Subramanian and Kapoor, 2013). Consistently, we found when treated with low concentrations of Docetaxel, the mitotic arrest was unable to sustain even in SAC proficient HeLa cells. Instead, the cells slip to the subsequent interphase without proper chromosome segregation and cytokinesis, yielding multinucleated cells. If the function of SAC is to protect against mis-segregation of single chromosome, then why didn’t a switch-like ‘all-or-none’ response evolve? (Subramanian and Kapoor, 2013) Here we hypothesize that the primary function of SAC is preservation of cell viability through avoiding mis-segregation of large numbers of rather than single chromosome. In support of this, the SAC is found to be essential for viability in vertebrates and SAC gene knockout mice are embryonic lethal with failure of chromosome segregation (Dobles et al., 2000).

The function of SAC to guard against CIN is irrelevant in tumor cells which are already aneuploidy. As a consequence of the abnormal number of chromosomes, tumor cells may require more time to properly align chromosomes at metaphase (Yang et al., 2008). Moreover, cancer cells frequently contain supernumerary centrosomes, which need to be clustered into two functional poles, thus facilitating a seemingly normal bipolar mitosis (Storchova and Kuffer, 2008). By delaying anaphase onset, SAC provide cancer cells with additional time to deal with these problems. Therefore, compared with non-transformed cells tumor cells exhibit a prolonged duration of mitosis and thus be more dependent on SAC. This explains why mutations in spindle checkpoint genes are rare and the mitotic checkpoint is not weakened but rather over-activated in many human tumors (Hernando et al., 2004). Notably, cancer cells were reported to be more sensitive to undergo cell death in response to checkpoint abrogation (Manchado et al., 2012).

Along this line, we propose that SAC plays a protective role when tumor cells are exposed to agents which potentially cause defects in chromosome alignment and segregation, such as taxanes. By blocking cell cycle progression, activation of SAC provides tumor cells with additional time to minimize errors in microtubule– kinetochore attachment and chromosome distribution, thereby increase the probability of cell survival. Accordingly, inhibition of SAC was reported to synergize with paclitaxel by elevating the frequency of chromosome mis-segregation (Janssen et al., 2009).

On the other hand, A compromised SAC has been associated with drug resistance. Tumor cells with compromised SAC tend to undergo mitotic slippage following taxane treatment, thus the cell death due to prolonged mitotic arrest is escaped. However, growing evidence suggest that cells following therapy can take a long time to die (Zasadil et al., 2014). We have demonstrated in this study that promotion of mitotic slippage only delays taxane induced cell death but does not lead to drug resistance. Moreover, apoptosis is not the only option through which taxanes exert their anti-tumor effect. Inhibition of apoptosis often leads to compensatory increase in cellular senescence. The extensive use of short-term cell death assays appears to be the primary reason for the incorrect correlation of SAC abrogation to taxanes resistance. By using clonogenic assay, we demonstrate the opposite, that inhibition of SAC markedly enhances the long-term effect of different microtubule toxins in both checkpoint proficient and less proficient tumor cells. Recently, the antitumor efficacy of Mps1 kinase inhibitors in combination with paclitaxel was validated by preclinical studies (Wengner et al., 2016). Together, the evidence provided by us and others suggest that this combination therapy represents a valuable mechanism that is expected to improve therapeutic efficacy of microtubule toxins.

## EXPERIMENTAL PROCEDURES

### Plasmid Construction

The coding sequence of Survivin and Bcl-xL were cloned from cDNA of HeLa cells by polymerase chain reaction. DNA encoding Bcl-2 was synthesized and base optimized without changing amino acids owing to a high GC content by Sangon Biotech Co., Ltd. These coding sequences were first cloned into pcDNA3.1 (Invitrogen), whereupon site-directed mutagenesis was performed to generate phosphomimetic or deficient mutants of each gene. The WT or mutant sequences were then cloned into the lentiviral vector pLVX-Puro (Clontech, Mountain View, CA) for lentivirus production. The luciferase gene used for the control was cloned from the pGL3-Basic Vector (Promega, Madison, WI). All inserts were confirmed by DNA sequencing. All primer sequences can be found in the Supplemental Information.

### Lentivirus Production

Lentiviral particles were produced according to the “pLKO.1 Protocol” provided by Addgene (Cambridge, MA). Briefly, WT and mutant sequences of Survivin, Bcl-2, and Bcl-xL were individually cloned into the pLVX-Puro vector. The resultant lentiviral vectors were co-transfected with the packaging plasmid psPAX2 (a gift from Didier Trono, Addgene plasmid # 12260) and envelope plasmid pMD2.G (a gift from Didier Trono, Addgene plasmid # 12259) into 293T cells using Lipofectamine 2000 (Invitrogen). After 12–15 h incubation, the medium was replaced with fresh DMEM + 10% FBS. Lentiviral particle-containing medium was harvested from cells after 48 h incubation and filtered through a 0.45-μm filter to remove the 293T cells, then was directly used to infect target cells.

### Cell Culture

HeLa, MDA-MB-231, MCF-7, and 293T cells were obtained from the Cell Bank of the Chinese Academy of Sciences (Shanghai, China). HeLa and 293T cells were maintained in Dulbecco’s modified Eagle’s medium (DMEM) (Macgene Technology Ltd., Beijing, China) supplemented with 10% fetal bovine serum (FBS) (Gibco BRL, Life Technologies, Grand Island, NY). MDA-MB-231 cells were grown in L-15 medium (Macgene Technology Ltd.) containing 10% FBS. MCF-7 cells were maintained in DMEM with 10% FBS and 10 μg/ml insulin.

Cell lines stably expressing WT and mutant forms of Survivin, Bcl-2, and Bcl-xL were generated by infection with the indicated lentiviral particles, generated as described above. After selection in medium containing 2 μg/ml puromycin for a week, the uninfected cells were no longer viable and overexpression of Survivin, Bcl-2, and Bcl-xL was confirmed by western blot. The expression of mutant protein was confirmed by cDNA pyrosequencing as described below.

### Chemicals

Following chemicals were used for cell treatment, either used alone or combined with other chemicals: Docetaxel (Sanofi-Aventis, Gentilli, France), purvalanol A (Tocris, Bristol, United Kingdom), Cisplatin (Sigma, St. Louis, MO), paclitaxel (MCE, Shanghai, China), nocodazole (MCE), AZ3146 (MCE), ABT-737 (MCE).

### Pyrosequencing

Total RNA was isolated from stably transduced cell lines using RNAiso Plus (TaKaRa, Dalian, China) and reverse-transcribed with a PrimeScript™ cDNA Synthesis Kit (TaKaRa) according to the manufacturer’s instructions. Pyrosequencing of each cDNA was performed by Sangon Biotech Co., Ltd. (Shanghai, China).

### Cell Synchronization and Mitotic Index Assessment

Cell cycle was blocked by exposure to 2 mM thymidine for 24 hr, and released by washing twice followed by incubation in thymidine-free medium containing 10% FBS. At the various time points following docetaxel treatment, Hoechst 33342 solution was added to the growth medium to obtain a final concentration of 1 μg/ml followed by incubation at 37°C for 20 min. Cells were then observed and photographed under a fluorescence microscopy (ZEISS Axio Vert.A1). Mitotic indexes were determined by counting the percentage of cells with spherical morphology and condensed chromatin.

### Small Interfering RNA Transfection

All siRNAs were synthesized by GenePharma Co., Ltd. (Shanghai, China). siRNA sequences from previous reports were used to target Survivin (Ana et al., 2003), Bcl-xL (Bruey et al., 2007), AURKB (Hauf et al., 2003), or Mad2 #1 (Collin et al., 2013) and #2 (Kabeche and Compton, 2012). siRNA transfection was performed using the Lipofectamine RNAIMAX reagent (Invitrogen) according to the manufacturer’s instructions. The final siRNA concentration was 40 nM unless stated otherwise. All siRNA sequences can be found in the Supplemental Information.

### Cell Viability Assay

A Cell Counting Kit-8 (CCK-8) (Dojindo, Kumamoto, Japan) was used to assess cell viability according to the manufacturer’s instructions. Cell viability was determined by measuring the absorbance at 450 nm using a microplate reader.

### FACS Apoptosis Assay

Annexin V-FITC/PI staining was performed on fresh cells according to the manufacturer’s specifications (Dojindo). The percentages of apoptotic cells were quantified by combining both early (Annexin V+/PI−) and late (Annexin V+/PI+) apoptotic cells.

### DNA content analysis

Cells following treatment were collected and fixed with 70% ethanol, and were subsequently incubated with 50 μg/ml propidium iodide (PI) with RNase A at 37°C for 30 min, and analyzed by FACS.

### Western blot analysis

Cell pellets were lysed in RIPA buffer (Macgene Technology Ltd., Beijing, China) with 10 μl/ml protease inhibitor cocktail (P8340, Sigma) and 10 μl/ml phosphatase inhibitor cocktail (p0044, Sigma). The proteins in cell lysates were separated by electrophoresis on a 15% SDS-polyacrylamide gel, transferred to nitrocellulose membranes, immunostained, and visualized by enhanced chemiluminescence detection reagents (Applygen Technologies Inc., Beijing, China). Antibodies against Survivin (#2808), Bcl-2 (#2870), Bcl-xL (#2764), Phospho-Histone H3 (#9713), phospho-Survivin (#8888), and phospho-Bcl-2 (#2827) were purchased from Cell Signaling (Danvers, MA), and the antibody specific for phospho-Ser-62-Bcl-xL was purchased from Abcam (Cambridge, UK). The β-Actin antibody was purchased from Zsbio (Beijing, China).

### Real-time RT-PCR

Total RNA was isolated using RNAiso Plus (TaKaRa), and reverse-transcribed using PrimeScriptTM RT reagent kit (TaKaRa) according to the manufacturer’s instruction. Real-time PCR was performed using SYBR premix EX Taq (TaKaRa) and analyzed with The CFX96 Touch System (Bio-Rad). Primers specific for GAPDH, Survivin, AURKB and Mad2 were designed and synthesized by TaKaRa. The data were represented as mean±SD from two independent experiments.

### SA-β-Gal assay

Cells were seeded in 12-well plates. Following indicated treatments, cells were allowed to grow in drug-free medium for more than 1 week. Senescent cells were detected by using Senescence β-Galactosidase Staining Kit (#9860 Cell Signaling) following the manufacturer’s instructions.

### Colony formation assays

Cells were seeded in 12-well plates. Following indicated treatments, cells were allowed to grow in drug-free medium for 2 weeks (HeLa) or 4 weeks (MDA-MB-231). And then fixed with methanol, and stained by crystal violet. The colony is defined to consist of at least 50 cells.

## SUPPLEMENTAL INFORMATION

Supplemental Information including all primer and siRNA sequences can be found along with this paper.

## AUTHOR CONTRIBUTIONS

T.L.H. designed the research, performed all experiments and wrote the manuscript.

Z.X.J. provided funding and laboratory.

Y.T.L., D.S.W., J.J., H.L. and B.H. supported the research.

H.S. secured funding, provided critical advice and supported the research in many aspects.

## ACKNOWLEDGMENTS

We thank Randy Schekman, Tony Hunter and the editors of eLife. Without their guidance this work cannot be. We thank Wei Huang (Tongji Medical College Huazhong University of Science and technology, Wuhan, China) for technical assistance in the preparation of this manuscript. No conflict of interest.

## SUPPLEMENTAL INFORMATION

**Figure S1.**
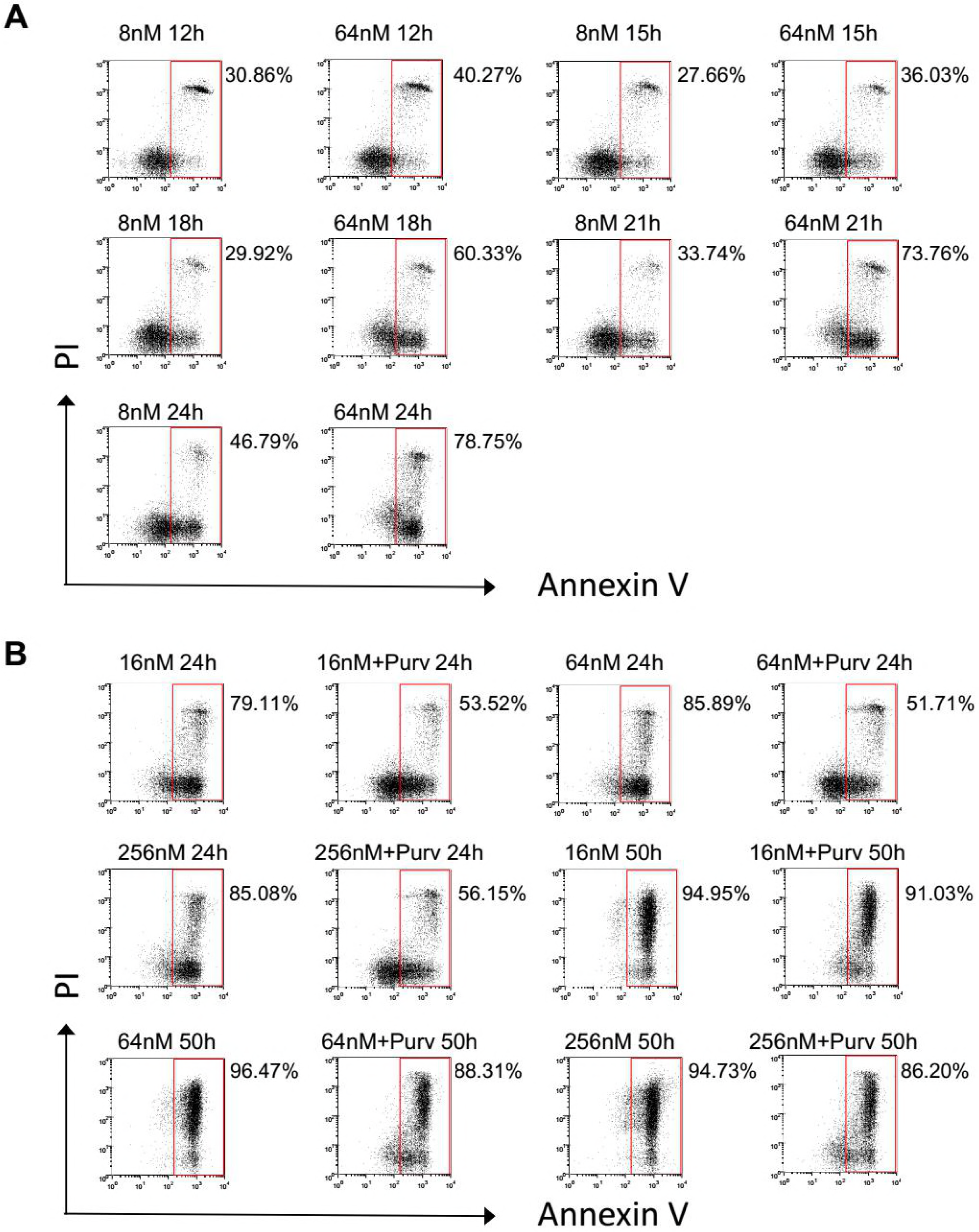
The relationship between mitotic arrest and treatment response to Docetaxel. (A) Synchronized HeLa cells were treated with different concentration of docetaxel and collected at indicated time intervals. Percentage of apoptosis were detected by FACS analysis following Annexin V/PI staining. (B) Synchronized cells were treated different concentrations of Docetaxel followed by addition of Purv, which facilitates mitotic slippage. Cells were collected after 24 hr or 50 hr Docetaxel treatment with or without Purv, and analyzed for induction of apoptosis.

**Figure S2.**
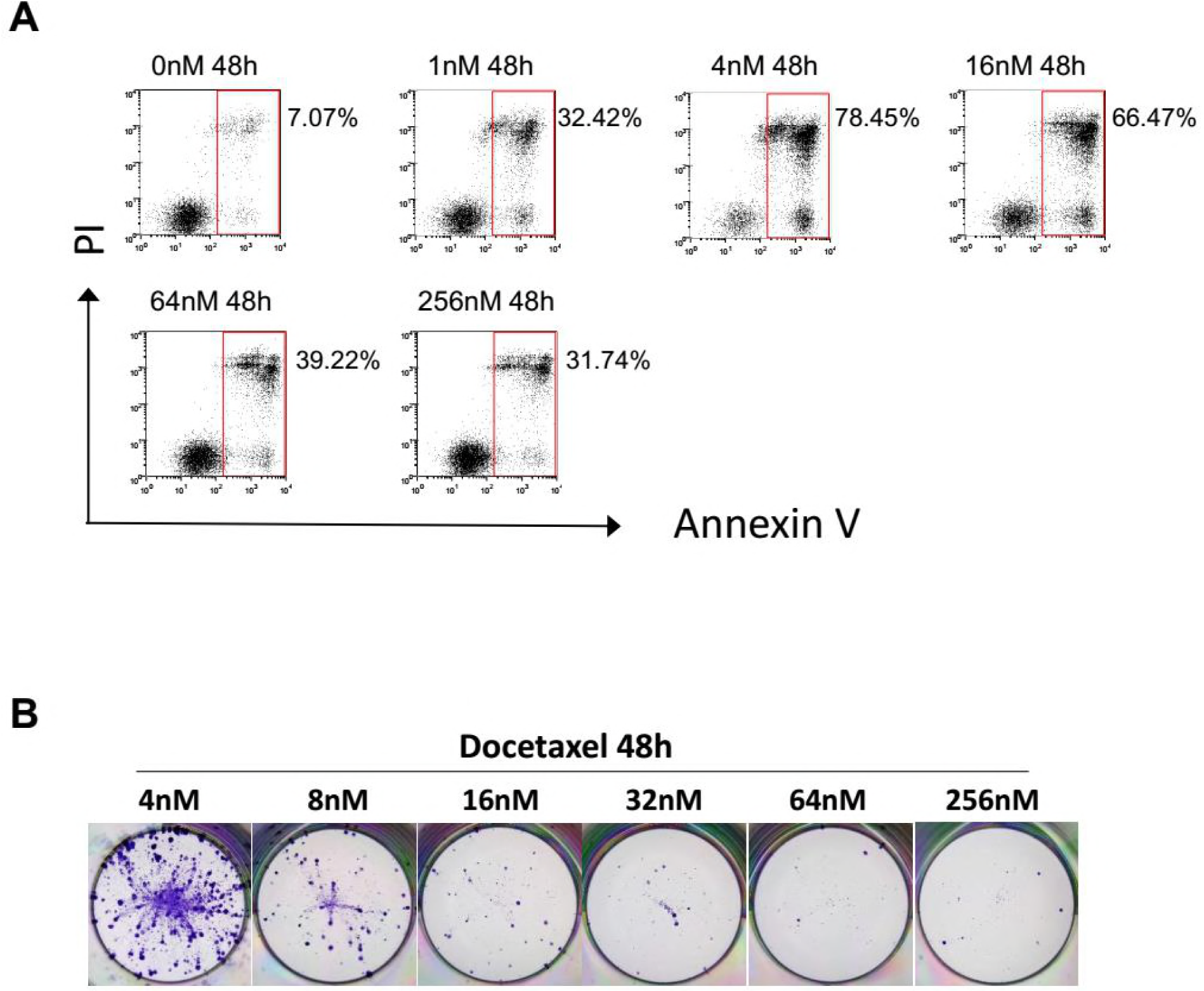
Apoptosis rates does not predict colony outgrowth after drug withdraw. (A) MDA-MB-231 cells were treated with the indicated concentrations of docetaxel for 48h, then collected and analyzed for induction of apoptosis. (B) Colony outgrowth assay. Cells were treated with the indicated concentrations of docetaxel for 48h, and then cultured in drug-free medium for 4 weeks. Colony outgrowth was photographed following fixation and staining with crystal violet.

**Figure S3.**
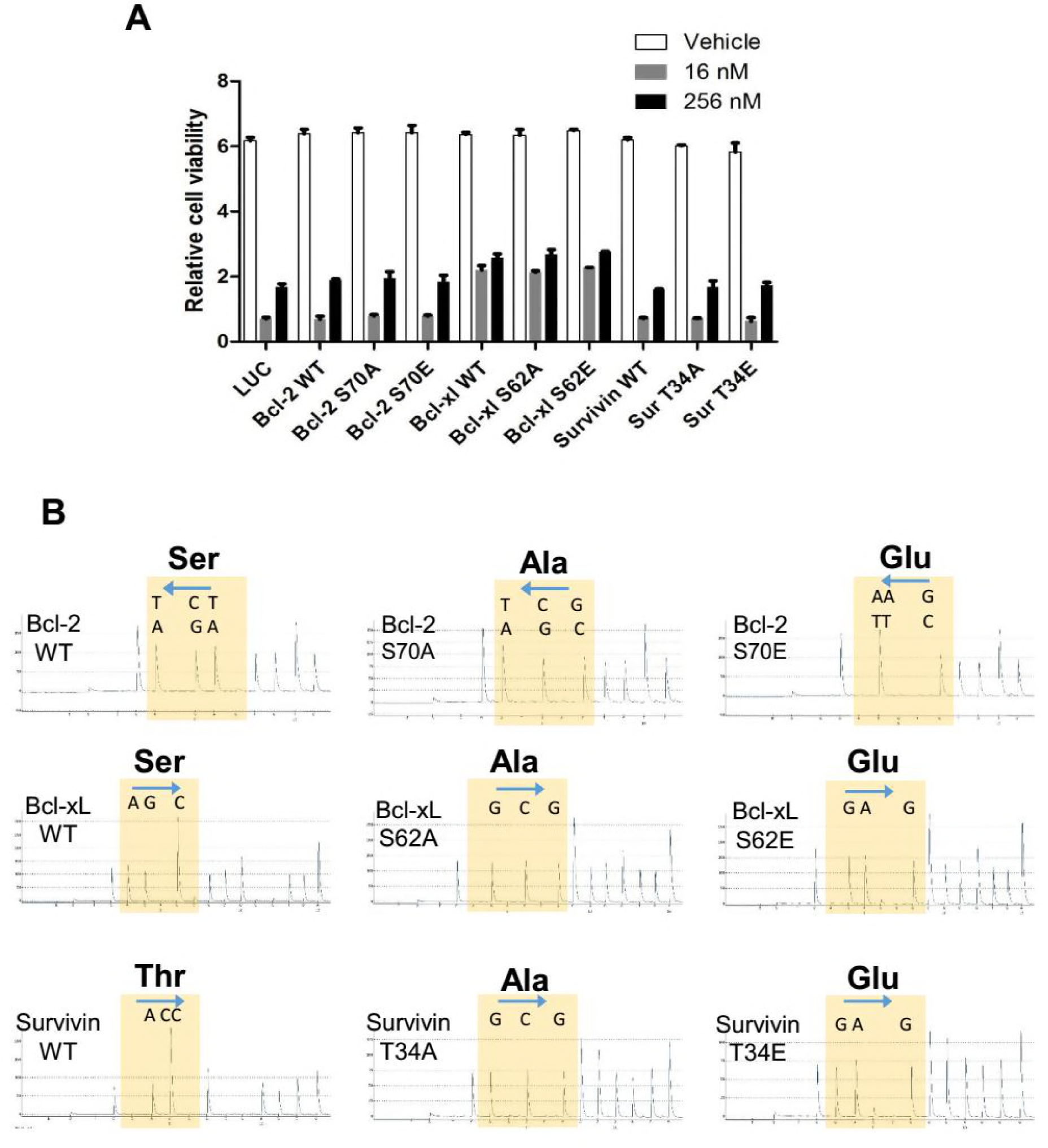
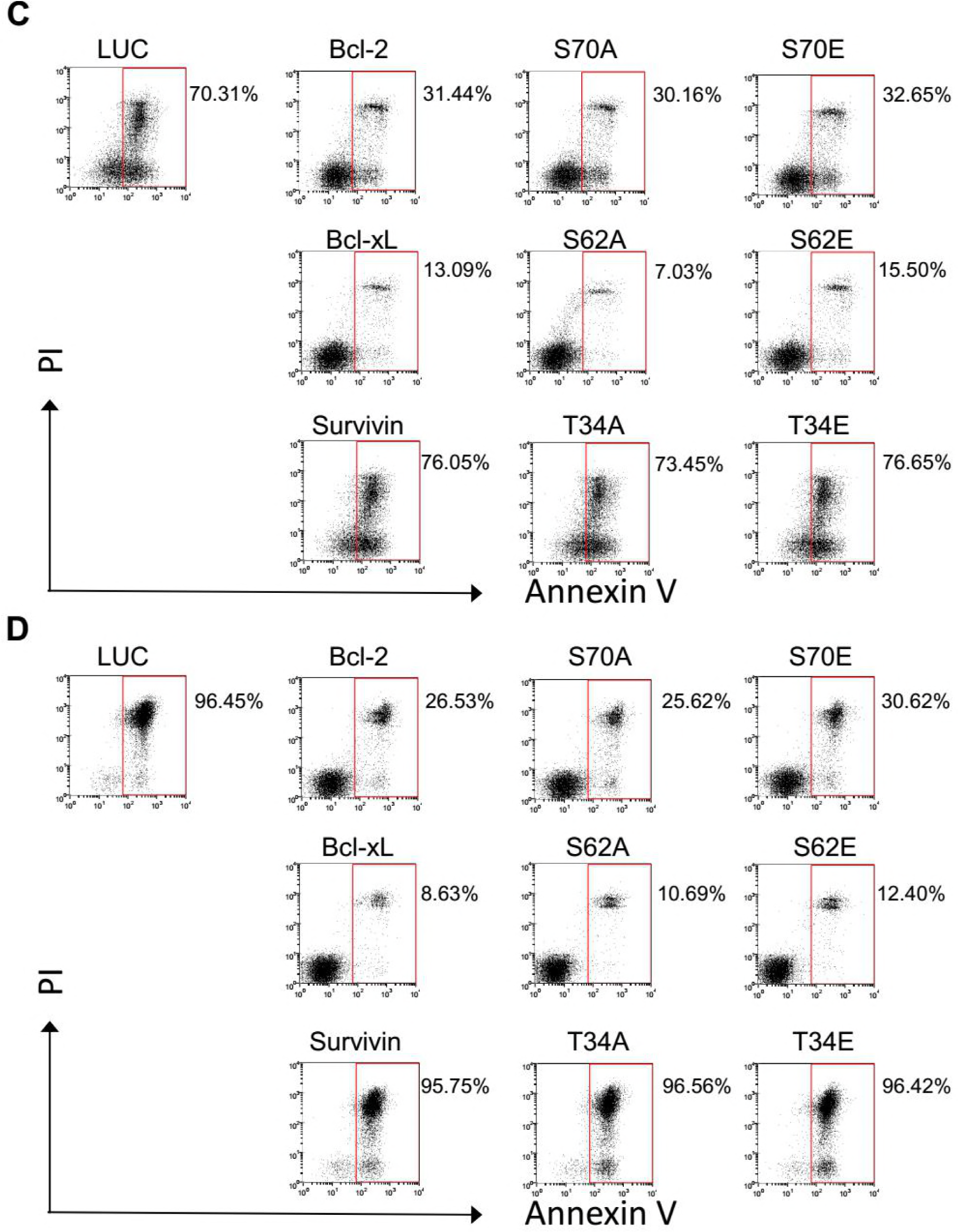
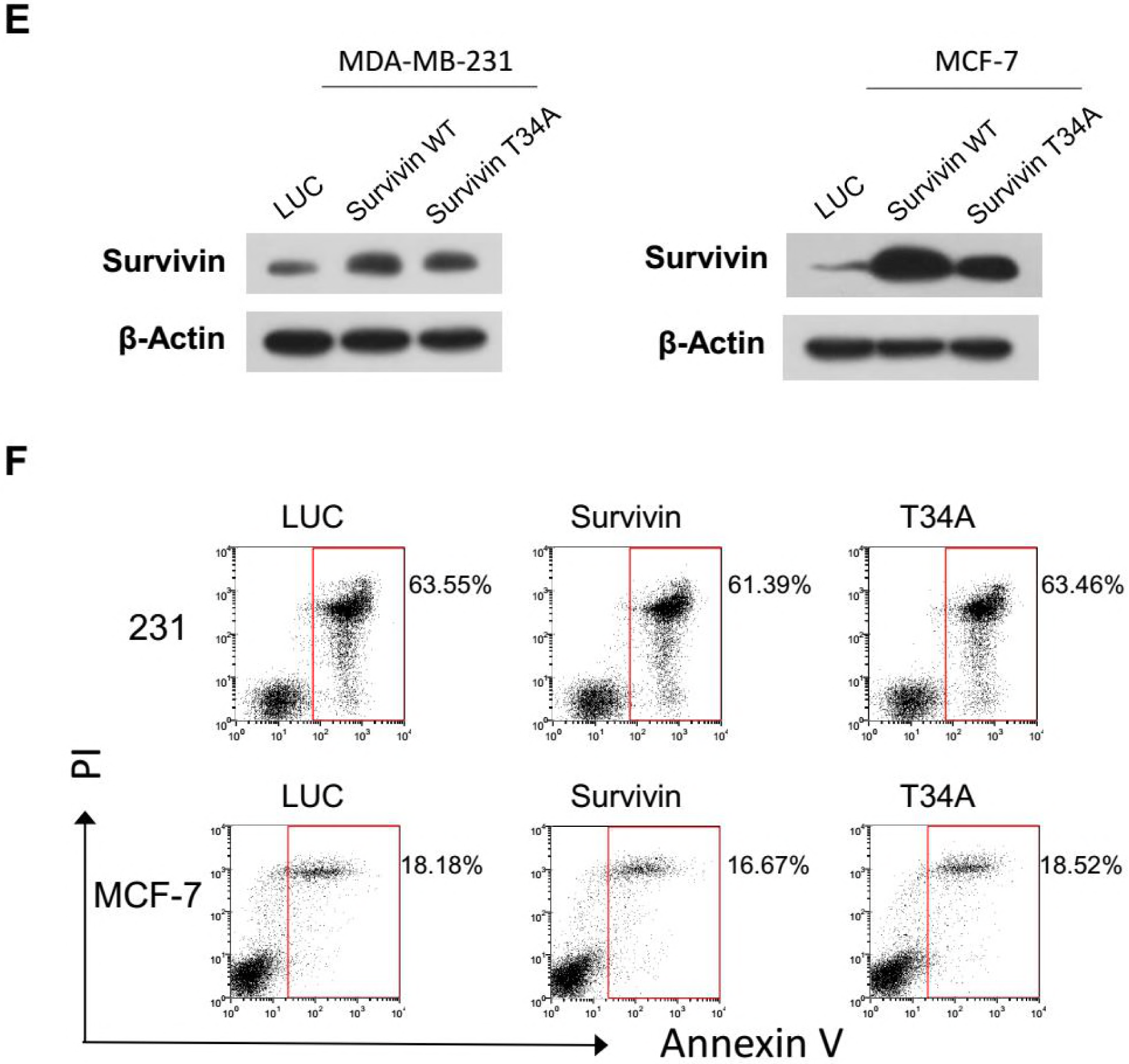
Overexpression of Bcl-2/Bcl-xL suppresses Docetaxel induced apoptosis but does not promote clonogenic survival. (A) MDA-MB-231 cells were infected with lentiviral vectors expressing the indicated proteins for 48 h, followed by docetaxel treatment for another 48 h. Lentiviral vectors expressing the luciferase (LUC) gene was used as a control. Cell viability was determined using the cck-8 reagent. Values represent the means ± S.D. (n = 2 wells). (B) Validation of mutant expression by cDNA pyrosequencing. Total RNA was isolated from stable cell lines and reverse-transcribed into cDNA. Pyrosequencing of each cDNA was performed by Sangon Biotech. (C and D) Different sensitivities of stable cell lines expressing Bcl-2, Bcl-x, or Survivin. Stable cell lines were treated with docetaxel (256 nM for 36 h) (C) or cisplatin (256μM for 36 h) (D). Cells were collected and stained for Annexin V/PI and analyzed by FACS. (E) Validation of Survivin expression in stable MDA-MB-231 or MCF-7 cell lines. Stable MDA-MB-231 or MCF-7 cell lines expressing LUC control, WT Survivin or T34A mutant were generated by lentiviral infection followed by selection with puromycin. The expression of the indicated protein was determined by western blotting. (F) Overexpression of WT Survivin or T34A mutant does not change the sensitivity of MDA-MB-231 or MCF-7 cells to Docetaxel. Stable cell lines were treated with docetaxel (16 nM for MDA-MB-231, 256 nM for MCF-7) for 72 hr, then the cells were collected, stained for Annexin V/PI, and analyzed by FACS.

**Figure S4.**
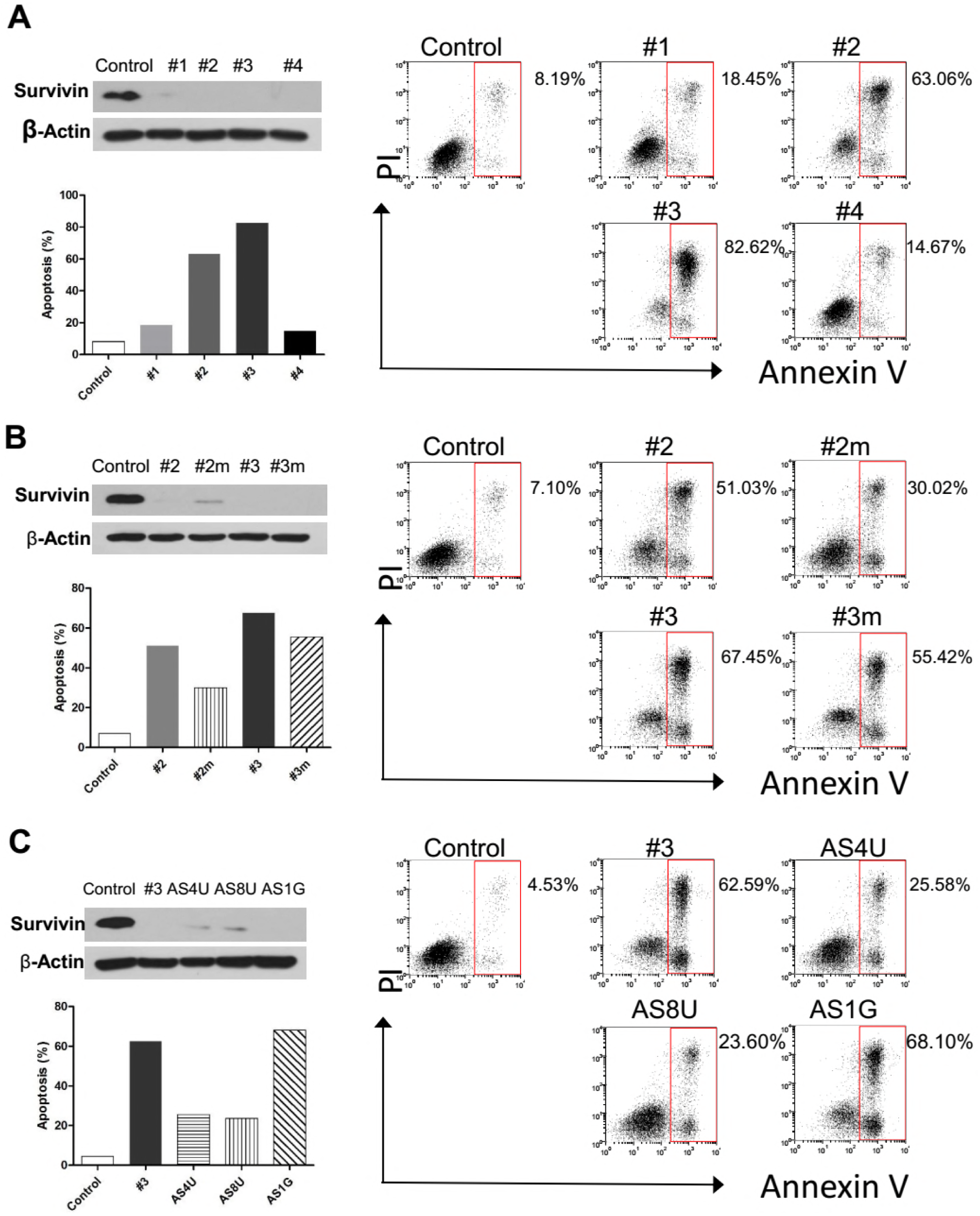
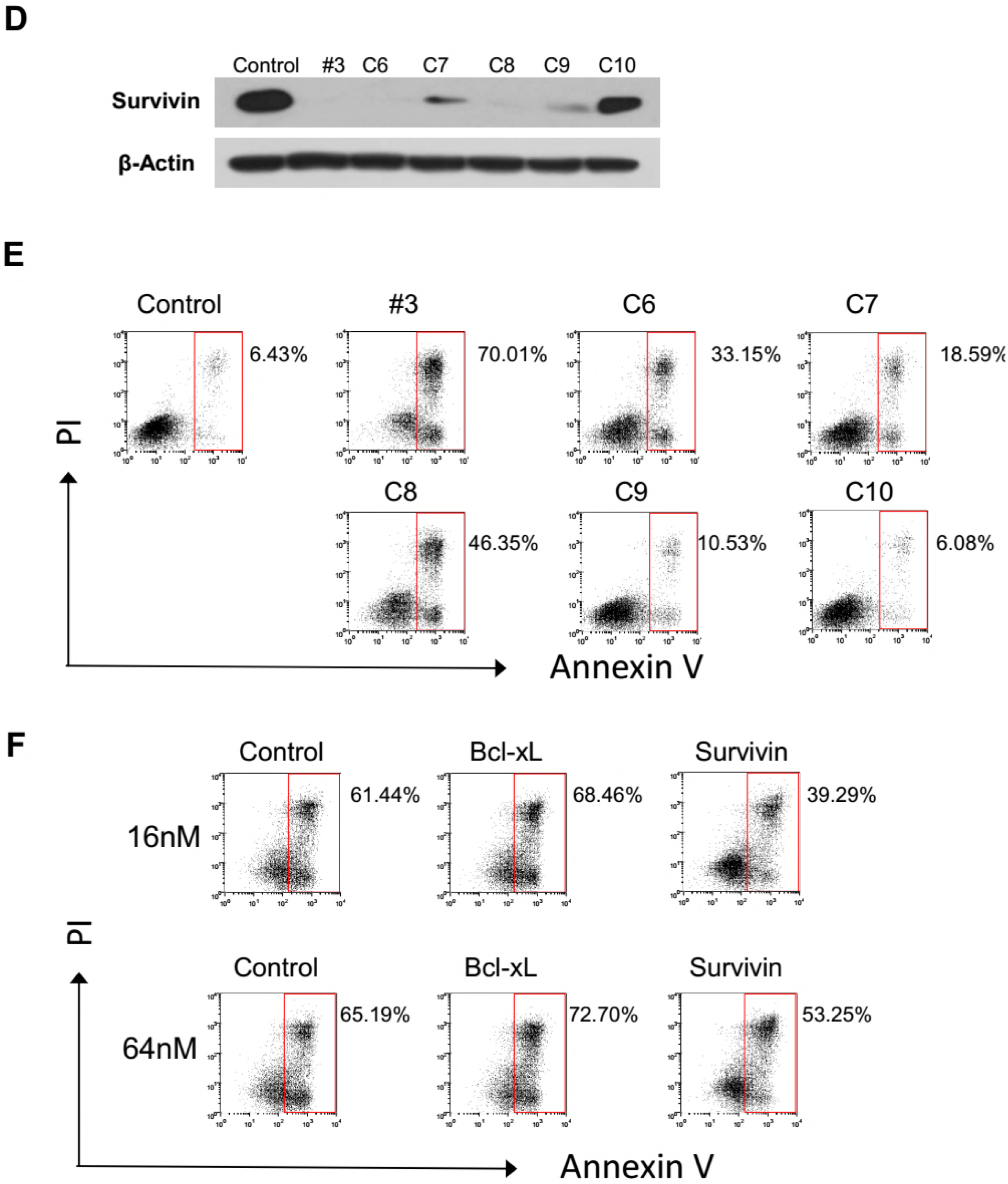
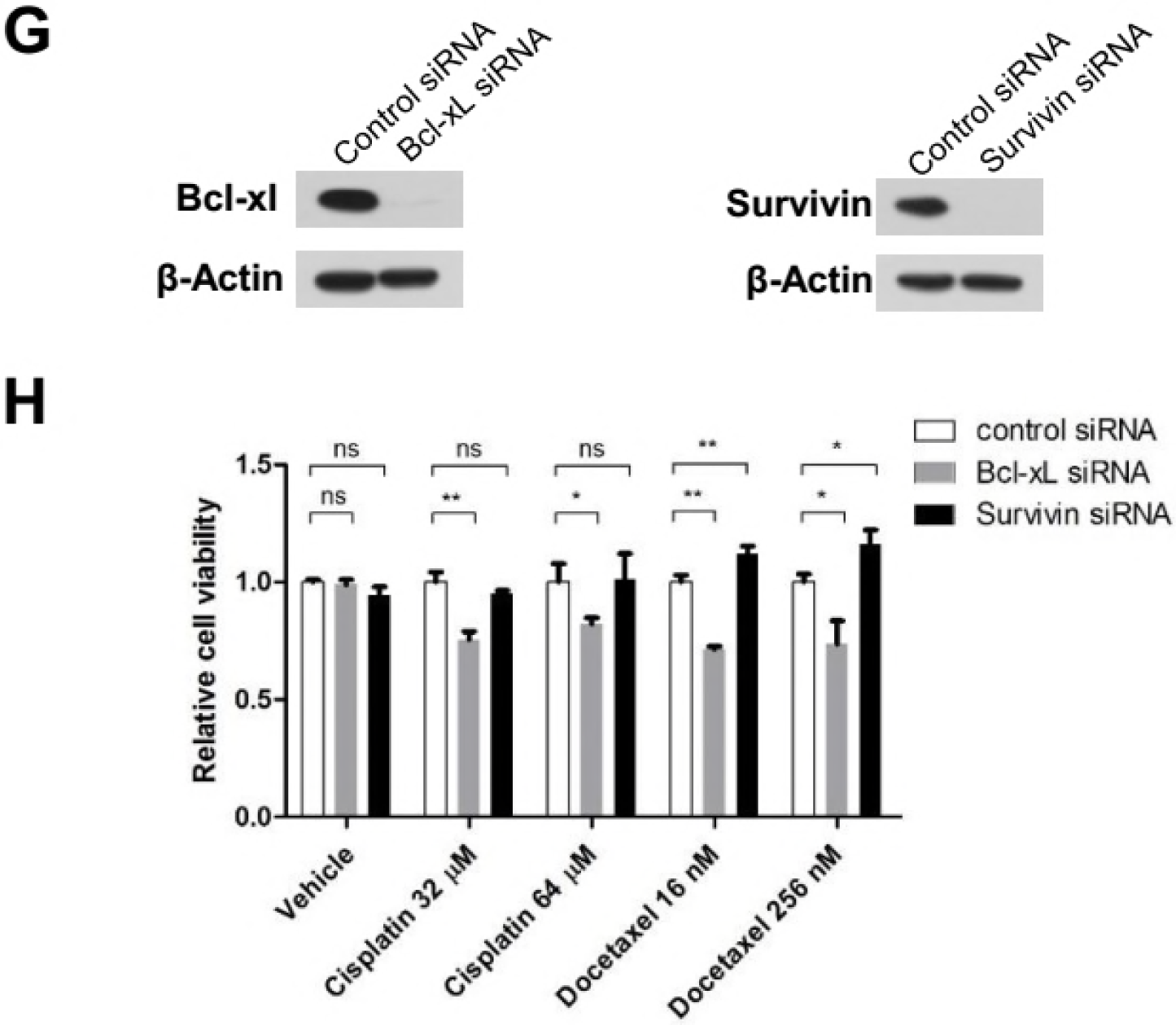
D Depletion of Survivin suppresses apoptosis induced by Docetaxel. (A) Varied levels of apoptosis induced by survivin siRNAs. 4 siRNA sequences were used to target Survivin: #1 was generated using siRNA WizardTM version 3.1 (InvivoGen, San Diego, CA); #2, #3 and the negative control siRNA were designed by GenePharma; the sequence of #4 was from a previous literature which exhibited minimal toxicity and was used for following experiments. HeLa cells incubated with the indicated siRNA for 48 h were harvested and analyzed for Survivin expression by western blotting. Apoptosis induced by individual Survivin siRNAs was determined by FACS analysis following Annexin V/PI staining. (B) Reduced toxicity of Survivin siRNA through chemically modification. The 2′-O-methyl modifications of the siRNAs were designed according to previous studies (Jackson et al., 2006). siRNAs #2m and #3m contained 2’-O-methyl modifications of positions 1 + 2 of the sense strand and position 2 of the guide strand. Cells were treated as described in (A). (C) Reduced toxicity of Survivin siRNA through antisense modification. The siRNA AS4U contained 2’-O-methyl modification of 4 uridine residues in the seed region of the antisense strand. The siRNA AS8U contained 2’-O-methyl modification of all 8 uridine residues in the antisense strand. The siRNA AS1G contained 2’-O-methyl modification of 1 guanosine residue in the antisense strand. Cells were treated as described in (A). (D) Depletion of Survivin by siRNAs with single base substitution. Mismatch designs C6-C10 were generated by replacing bases 6–10 with the complement of the original siRNA, respectively. HeLa cells incubated with the indicated siRNA for 48 h were harvested and analyzed for Survivin expression by western blotting. (E) Apoptosis induced by siRNAs with single base substitution was determined by FACS analysis following Annexin V/PI staining. (F) HeLa cells were incubated with the indicated siRNAs for 36 hr followed by 16nM or 64nM Docetaxel treatment for 24h. Cells were collected and analyzed for induction of apoptosis. (G) Depletion of Bcl-xl and Survivin in MCF-7 cells. MCF-7 cells incubated with the indicated siRNAs for 48 h were harvested and analyzed for Survivin expression by western blotting. (H) MCF-7 cells were incubated with the indicated siRNAs for 24 h followed by docetaxel or cisplatin treatment for 6 days. Cell viability was determined using the cck-8 reagent. Values represent the means ± S.D. (n = 3 wells). **p < 0.01; *p < 0.05; ns, not significant. Unpaired and two-tailed t test was used.

**Figure S5.**
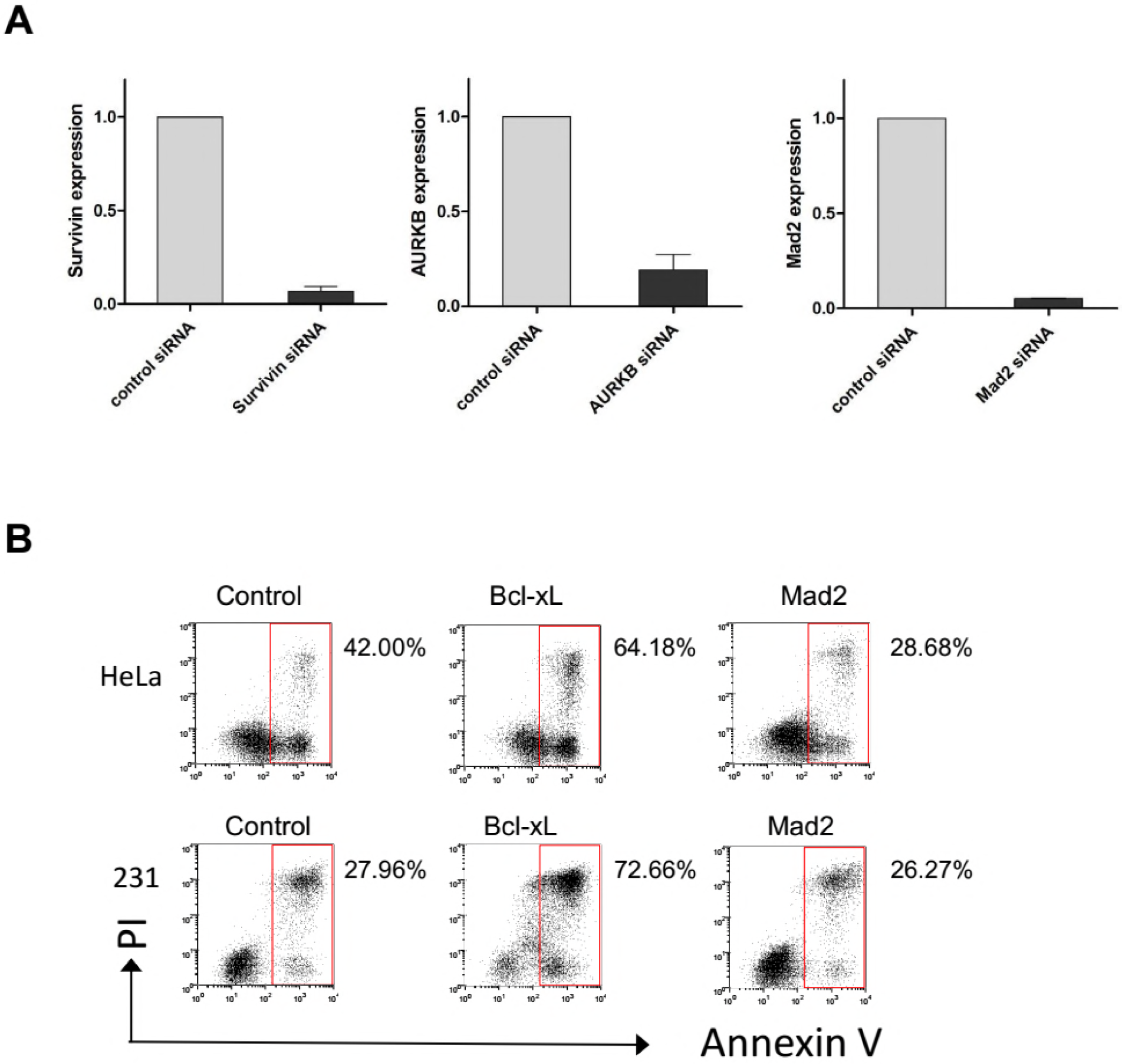
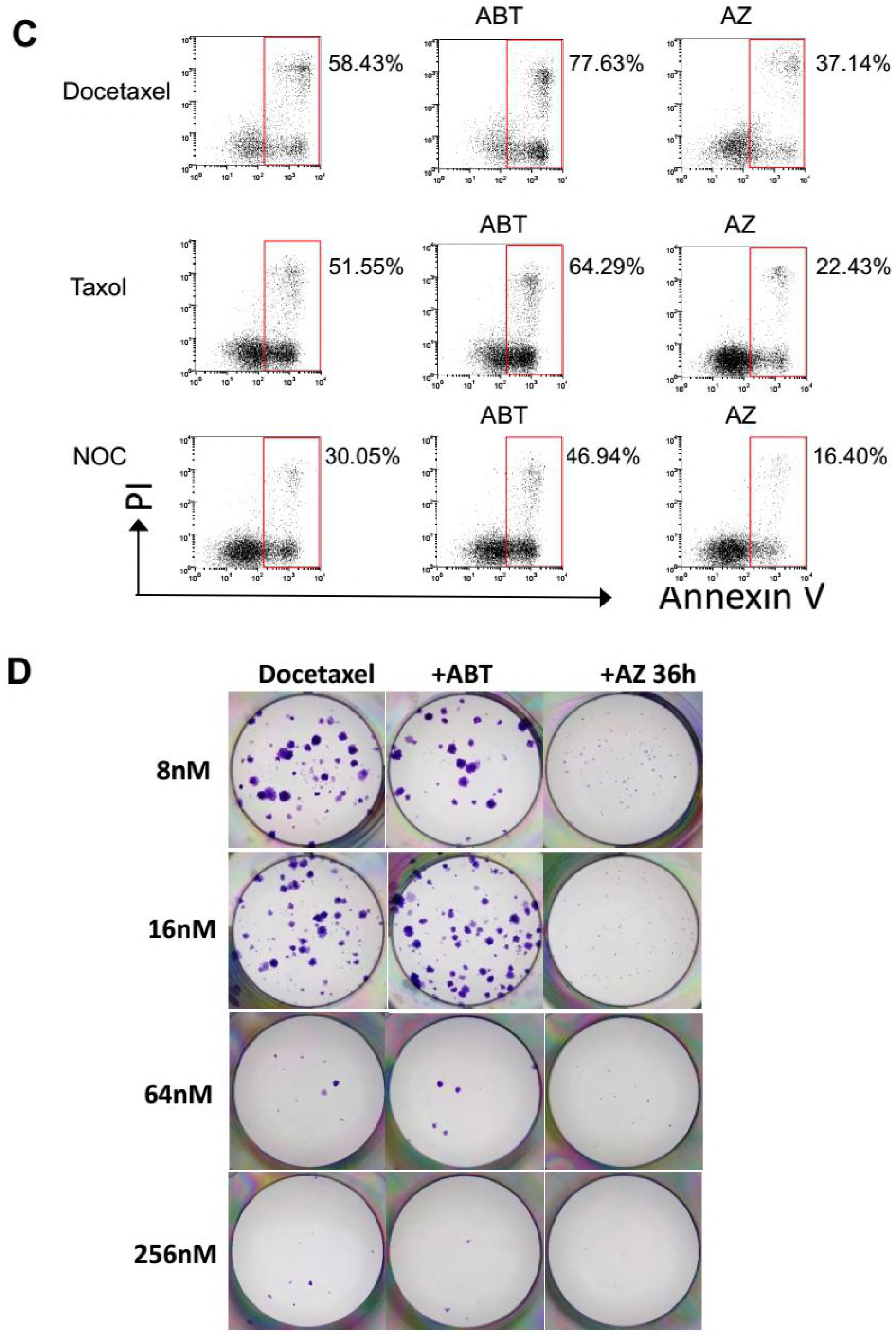
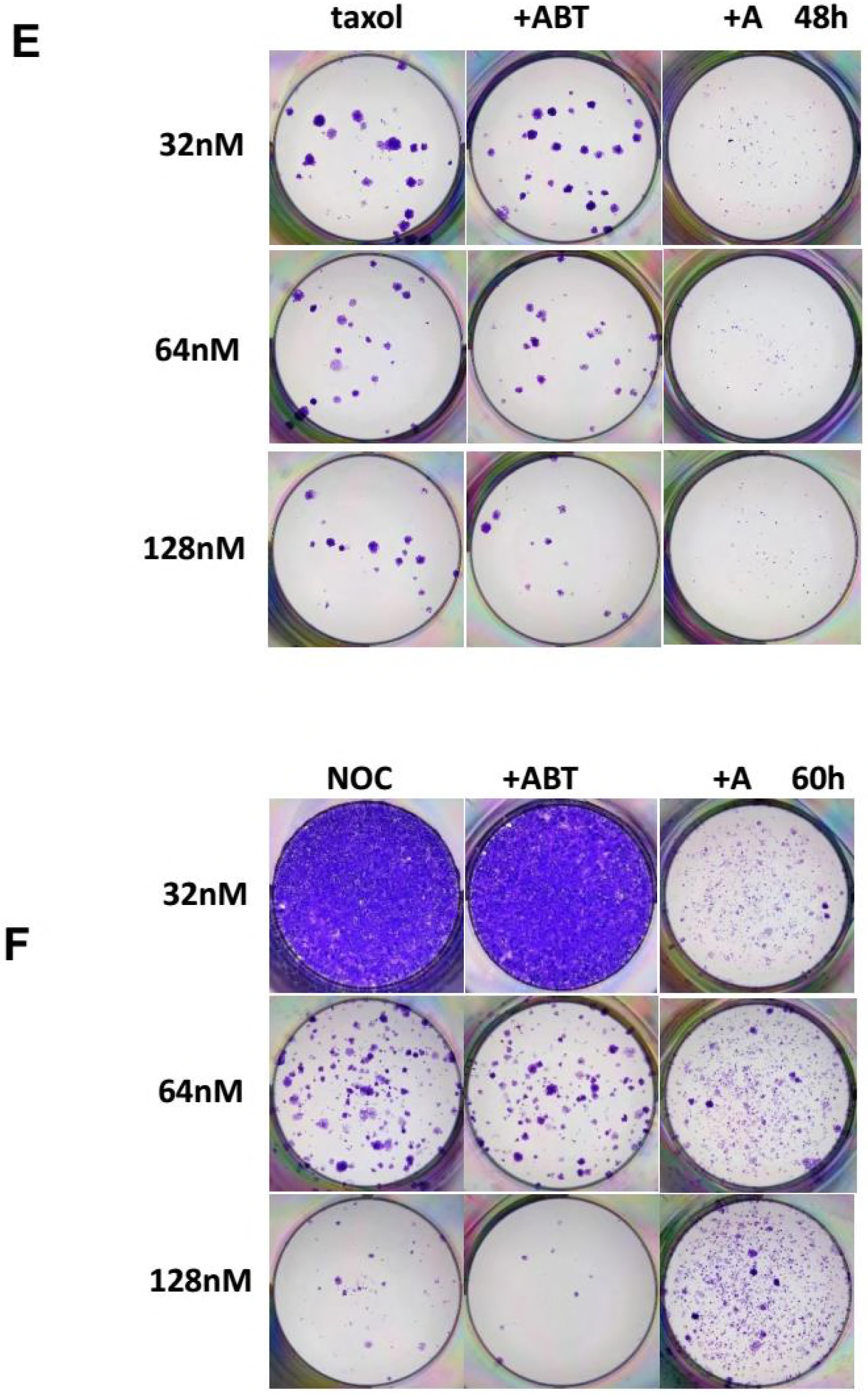
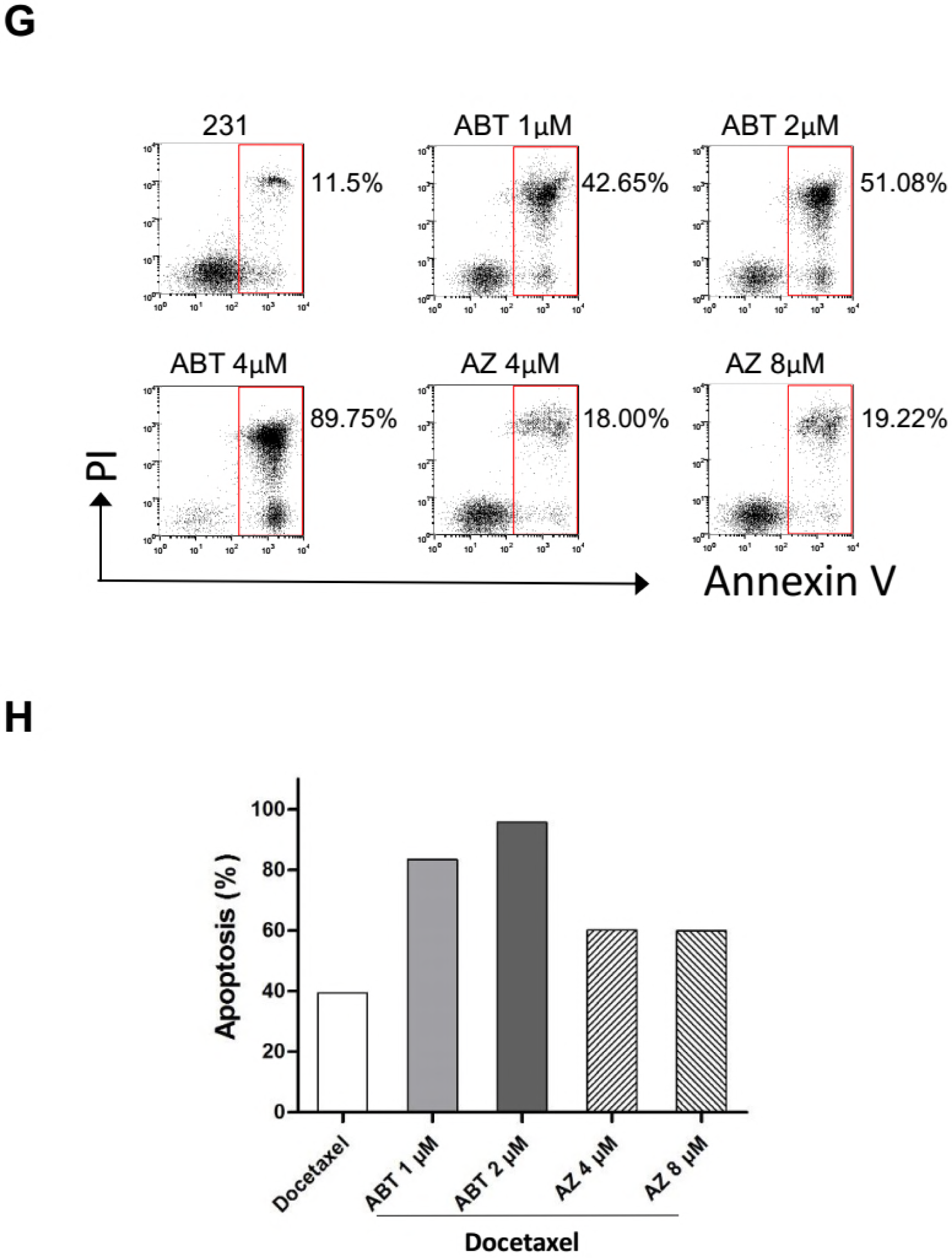
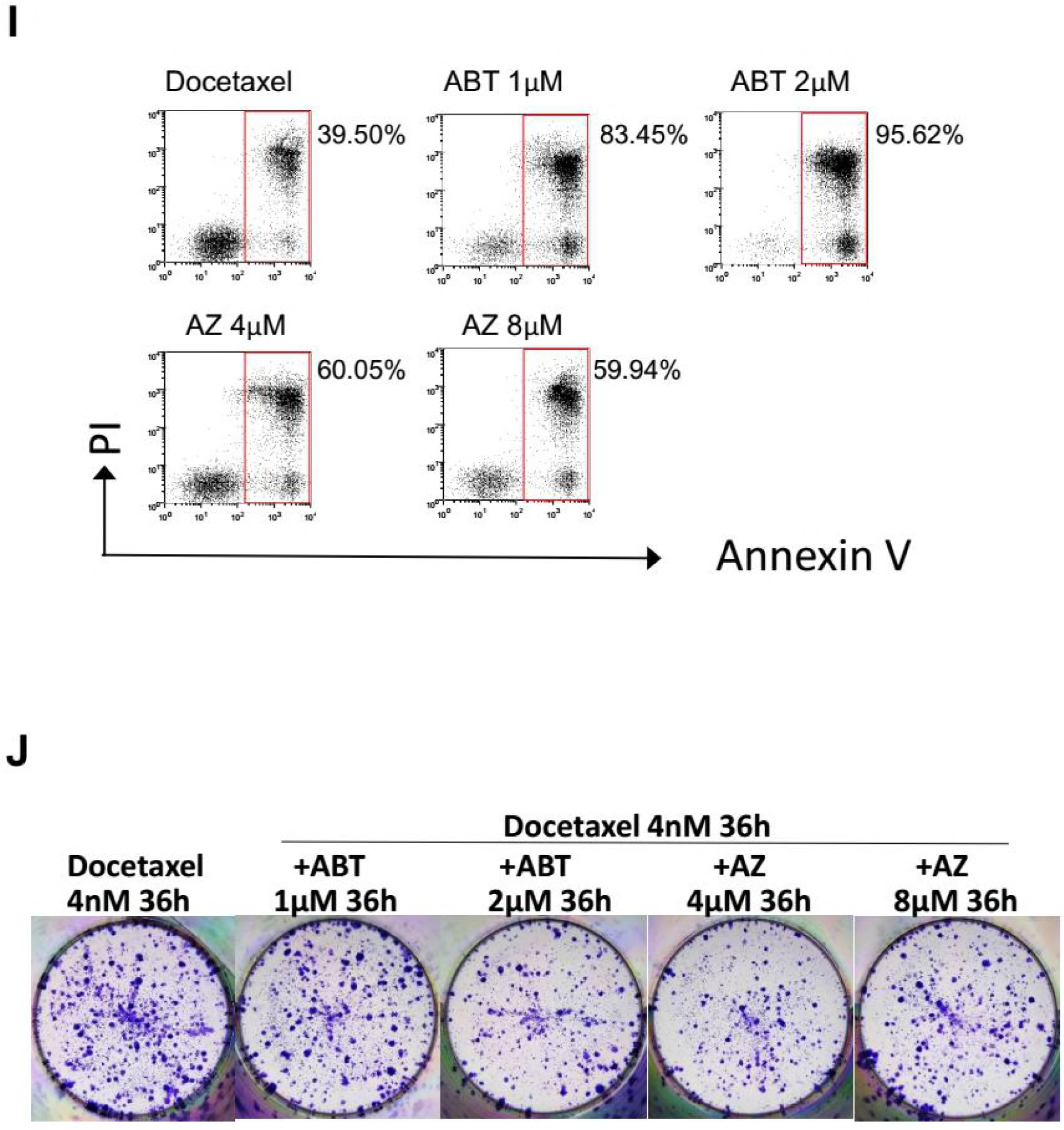
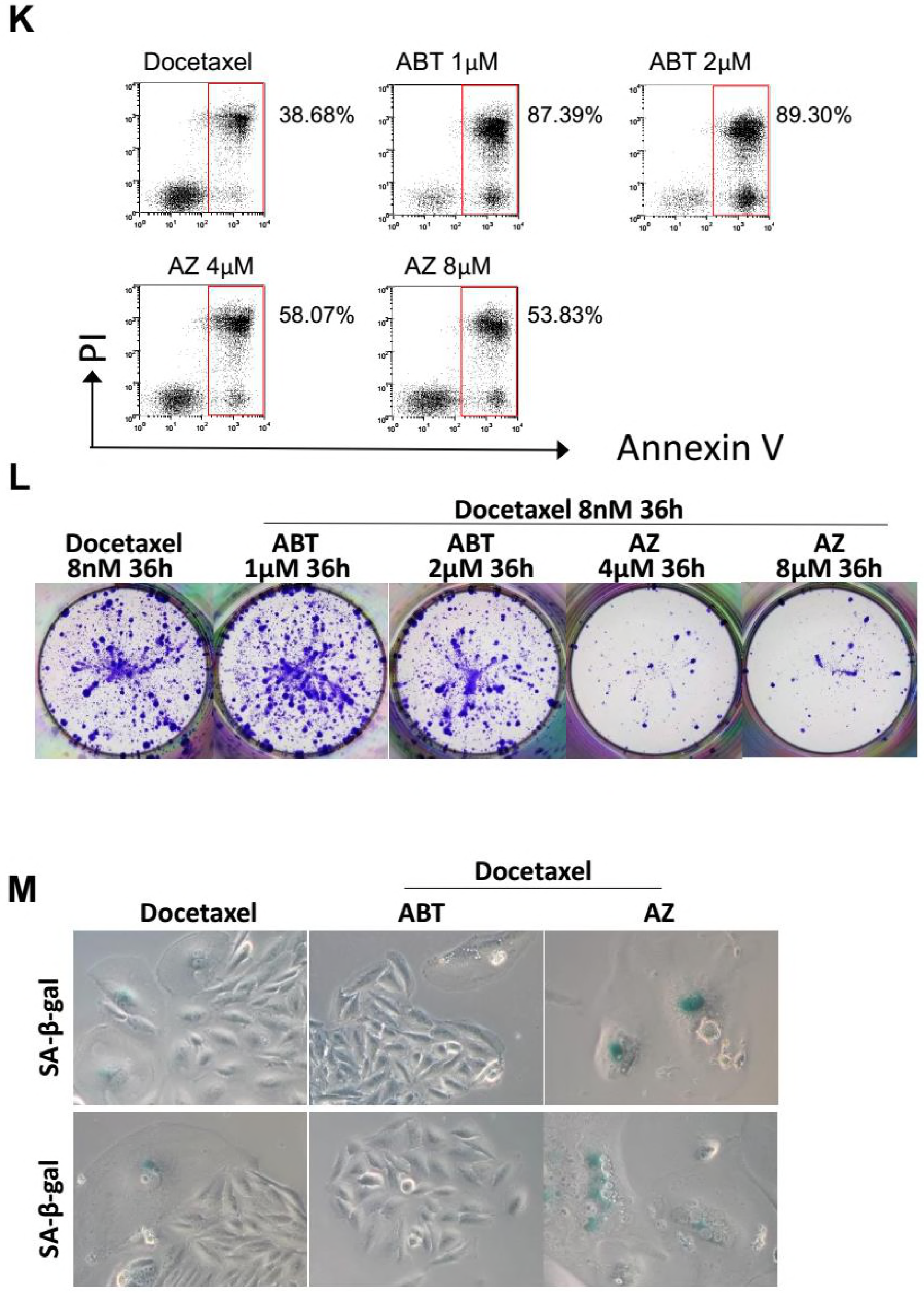
Inhibition of the SAC suppresses apoptosis induced by microtubule toxins but enhances their long-term efficacy. (A) HeLa cells incubated with the indicated siRNA for 48 hr were harvested and subjected to quantitative real-time PCR analysis for expression of Survivin, AURKB and Mad2, respectively. Values represent mean ± s.d. (n = 2 independent experiments). (B) HeLa and MDA-MB-231 cells were incubated with indicated siRNAs for 48h and then exposed to Docetaxel for 36h, cells were analyzed for induction of apoptosis. (C) HeLa cells were treated with microtubule toxins, including Docetaxel, paclitaxel or nocodazole, alone or in combination with ABT-737 or AZ3146, and then analyzed for induction of apoptosis. (D, E and F) HeLa cells were treated with the indicated concentrations of microtubule toxins, including Docetaxel (D), paclitaxel (E) or nocodazole (F), alone or in combination with ABT-737 (4μM) or AZ3146 (4μM), and then cultured in drug-free medium to allow for colony outgrowth. (G) MDA-MB-231 cells were treated with the indicated concentrations of ABT-737 or AZ3146 for 36h, then collected and analyzed for induction of apoptosis. (H, I and J) MDA-MB-231 cells were treated with the indicated concentrations of ABT-737 or AZ3146 in combination with Docetaxel for 36h. Cells were either collected and analyzed for induction of apoptosis (H and I), or cultured in drug-free medium to allow for Colony outgrowth (J). (K and L) MDA-MB-231 cells were first treated with the indicated concentrations of ABT-737 or AZ3146 for 24h, and then with ABT-737 or AZ3146 in combination with Docetaxel (8nM) for 36h. Cells were either collected and analyzed for induction of apoptosis (K), or cultured in drug-free medium to allow for Colony outgrowth (L). (M) HeLa cells were treated with Docetaxel, alone or in combination with ABT-737 or AZ3146. Cellular senescence was determined by SA-β-Gal staining.

**Figure S6.**
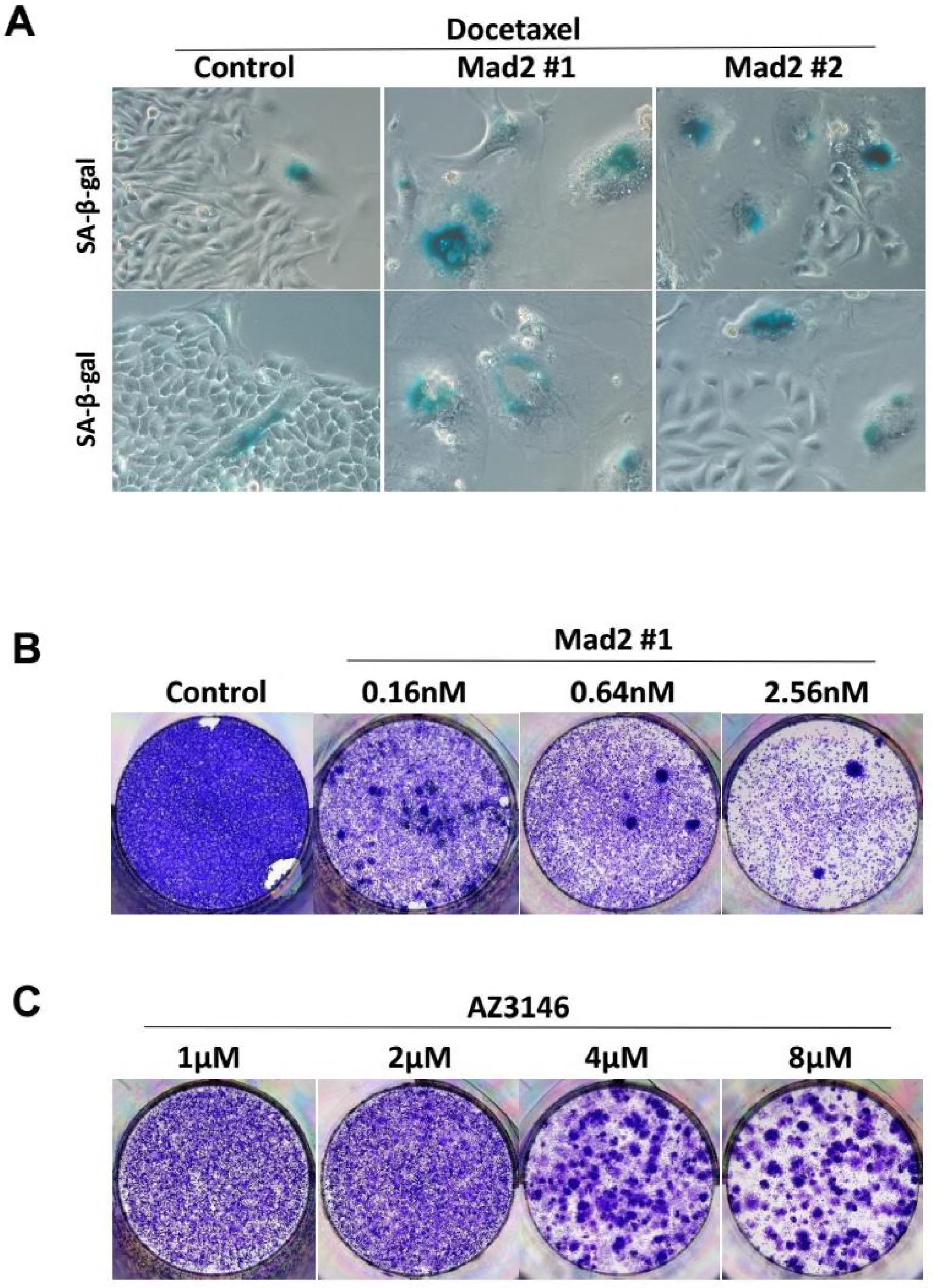
Partial Abrogation of SAC synergizes with Docetaxel in blocking colony formation. (A) HeLa cells were incubated with the indicated siRNAs for 48 hr followed by Docetaxel treatment for 36h, and then cultured in drug-free medium. Cellular senescence was determined by SA-β-Gal staining. (B) HeLa cells were incubated with the indicated concentrations of siRNAs for 48 hr, and then cultured in drug-free medium for 2 weeks and colony formation was photographed following fixation and staining with crystal violet. (C) HeLa cells were treated with the indicated concentrations of AZ3146 for 48 hr, and then cultured in drug-free medium for 2 weeks and colony formation was photographed following fixation and staining with crystal violet.

**Table S1.**
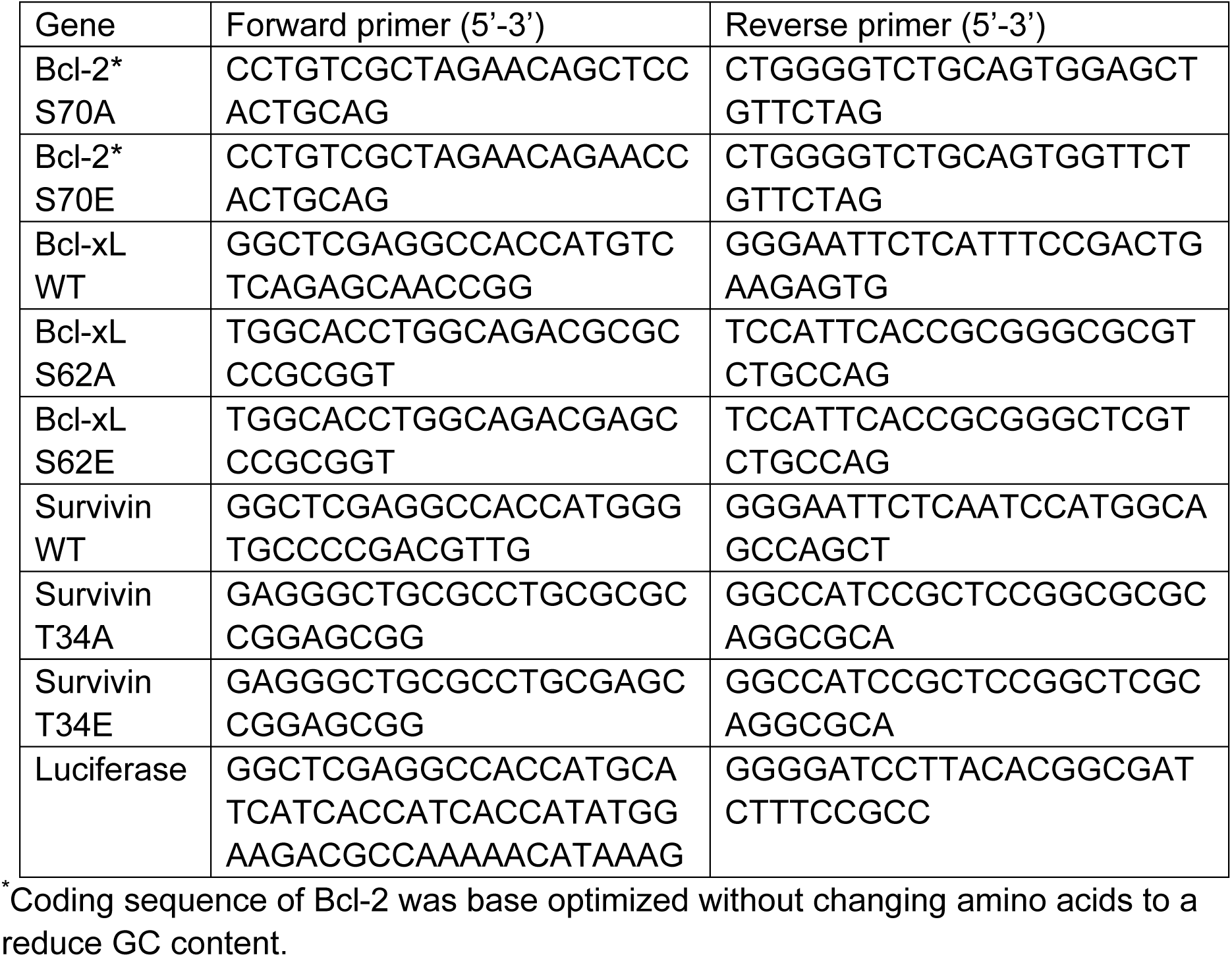
Sequences of PCR primers

**Table S2.**
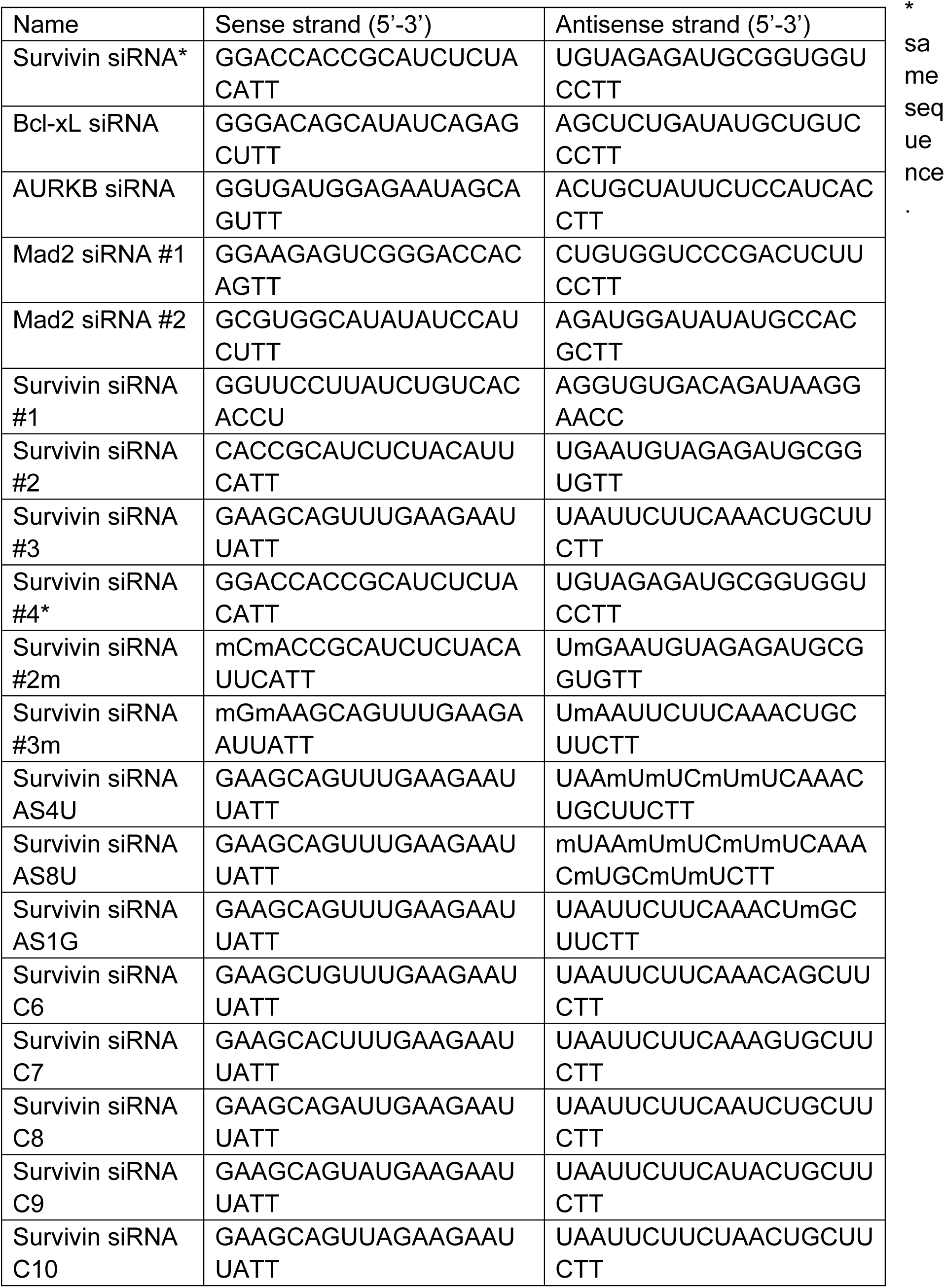
Sequences of siRNAs

